# Allosteric stabilization of calcium and phosphoinositide dual binding engages three synaptotagmins in fast exocytosis

**DOI:** 10.1101/2021.10.22.465434

**Authors:** Janus R. L. Kobbersmed, Manon M. M. Berns, Susanne Ditlevsen, Jakob B. Sørensen, Alexander M. Walter

## Abstract

Synaptic communication relies on the fusion of synaptic vesicles with the plasma membrane, which leads to neurotransmitter release. This exocytosis is triggered by brief and local elevations of intracellular Ca^2+^ with remarkably high sensitivity. How this is molecularly achieved is unknown. While synaptotagmins confer the Ca^2+^ sensitivity of neurotransmitter exocytosis, biochemical measurements reported Ca^2+^ affinities too low to account for synaptic function. However, synaptotagmin’s Ca^2+^ affinity increases upon binding the plasma membrane phospholipid PI(4,5)P_2_ and, vice versa, Ca^2+^-binding increases synaptotagmin’s PI(4,5)P_2_ affinity, indicating a stabilization of the Ca^2+^/PI(4,5)P_2_ dual-bound syt. We here devise a molecular exocytosis model based on this positive allosteric stabilization and the assumptions that (1.) synaptotagmin Ca^2+^/PI(4,5)P_2_ dual binding lowers the energy barrier for vesicle fusion and that (2.) the effect of multiple synaptotagmins on the energy barrier is additive. The model, which relies on biochemically measured Ca^2+^/PI(4,5)P_2_ affinities and protein copy numbers, reproduced the steep Ca^2+^ dependency of neurotransmitter release. Our results indicate that each synaptotagmin dual binding Ca^2+^/PI(4,5)P_2_ lowers the energy barrier for vesicle fusion by 4.85 k_B_T and that allosteric stabilization of this state enables the synchronized engagement of three synaptotagmins for fast exocytosis. Furthermore, we show that mutations altering synaptotagmin’s allosteric properties may show dominant-negative effects, even though synaptotagmins act independently on the energy barrier, and that dynamic changes of local PI(4,5)P_2_ (e.g. upon vesicle movement) dramatically impact synaptic responses. We conclude that allosterically stabilized Ca^2+^/PI(4,5)P_2_ dual binding enables synaptotagmins to exert their coordinated function in neurotransmission.

## Introduction

Regulated neurotransmitter (NT) release from presynaptic terminals is crucial for information transfer across chemical synapses. NT release is triggered by action potentials (APs), which are transient de- and repolarizations of the presynaptic membrane potential that induce Ca^2+^ influx through voltage-gated channels. The resulting brief and local elevations of the intracellular Ca^2+^ concentration ([Ca^2+^]_i_) trigger the fusion of NT-containing synaptic vesicles (SVs) from the so-called readily releasable pool (RRP), whose SVs are localized (docked) at the plasma membrane and molecularly matured (primed) for fusion (Kaeser and Regehr, 2017; Sudhof, 2013; Verhage and Sørensen, 2008). A high Ca^2+^ sensitivity of NT release is needed to achieve fast responses to the very short AP-induced Ca^2+^ transient and correspondingly, the SV fusion rate depends on the [Ca^2+^]_i_ to the 4^th^-5^th^ power (Bollmann et al., 2000; Burgalossi et al., 2010; Heidelberger et al., 1994; Schneggenburger and Neher, 2000). Accordingly, previous models of NT release have assumed the successive binding of five Ca^2+^ ions to a sensor that regulates release (Bollmann *et al*., 2000; Lou et al., 2005; Schneggenburger and Neher, 2000). However, how these macroscopic properties arise from the molecular components involved in SV fusion is still unknown.

The energy for SV fusion is provided by the assembly of the neuronal SNARE complex, which consists of vesicular synaptobrevin/VAMP and plasma membrane bound SNAP25 and syntaxin proteins (Jahn and Fasshauer, 2012; Sudhof, 2013). Ca^2+^ sensitivity of SV fusion is conferred by the vesicular protein synaptotagmin (syt), which interacts with the SNAREs (Brewer et al., 2015; Littleton et al., 1993; Mohrmann et al., 2013; Schupp et al., 2016; Zhou et al., 2015; Zhou et al., 2017). Several syt isoforms are expressed in synapses. Depending on the synapse type (e.g. mouse hippocampal pyramidal neurons or the Calyx of Held), syt1 or syt2 is required for synchronous, Ca^2+^-induced fusion (Geppert et al., 1994; Kochubey et al., 2016; Kochubey and Schneggenburger, 2011; Sudhof, 2013). These two syt isoforms are highly homologous and contain two cytosolic, Ca^2+^-binding domains, C2A and C2B (Sudhof, 2002), of which the C2B domain has been shown to be essential, and in some cases even sufficient, for synchronous NT release (Bacaj et al., 2013; Gruget et al., 2020; Kochubey and Schneggenburger, 2011; Lee et al., 2013; Mackler et al., 2002). The C2B domain contains two Ca^2+^ binding sites on its top loops (Fernandez et al., 2001). In addition, a second binding site allows the C2B domain to bind to the signaling lipid phosphatidylinositol 4,5-phosphate (PI(4,5)P_2_) located in the plasma membrane (Bai et al., 2004; Fernandez-Chacon et al., 2001; Honigmann et al., 2013; Li et al., 2006; Xue et al., 2008), but might also participate in (possibly transient) SNARE interactions (Brewer *et al*., 2015; Zhou *et al*., 2015; Zhou *et al*., 2017). A third site, located in the far end of the C2B domain (R398 and R399 in mouse syt1), is also involved in both SNARE- and membrane contacts (Nyenhuis et al., 2021; Xue *et al*., 2008; Zhou *et al*., 2015). Via these interactions, the syt C2B domain can induce close membrane-membrane contact *in vitro* (Araç et al., 2006; Chang et al., 2018; Honigmann *et al*., 2013; Nyenhuis *et al*., 2021; Seven et al., 2013; Xue *et al*., 2008), stable vesicle-membrane docking (Chang *et al*., 2018; Chen et al., 2021; de Wit et al., 2009), as well as dynamic vesicle-membrane association upon Ca^2+^ influx in the cell (Chang *et al*., 2018).

Despite its central role as the Ca^2+^ sensor for NT release, the intrinsic Ca^2+^ affinity of the isolated syt C2B domain is remarkably low (KD ≈ 200 µM, (Radhakrishnan et al., 2009; van den Bogaart et al., 2012)), much lower than the Ca^2+^ sensitivity of NT release (Bollmann *et al*., 2000; Schneggenburger and Neher, 2000). However, binding of the C2B domain to PI(4,5)P_2_, which is enriched at synapses (van den Bogaart et al., 2011a), drastically increases its Ca^2+^ affinity (van den Bogaart *et al*., 2012). Similarly, the affinity for PI(4,5)P_2_ increases upon Ca^2+^ binding (Pérez-Lara et al., 2016; van den Bogaart *et al*., 2012). This indicates a positive allosteric coupling between the binding sites for Ca^2+^ and PI(4,5)P_2_, which promotes dual binding of Ca^2+^/PI(4,5)P_2_ (Li *et al*., 2006; Radhakrishnan *et al*., 2009; van den Bogaart *et al*., 2012). As binding of both molecules to syt is involved in fusion (Kedar et al., 2015; Li *et al*., 2006; Mackler *et al*., 2002; Mackler and Reist, 2001; Wang et al., 2016; Wu et al., 2021a; Wu et al., 2021b), this positive allosteric coupling might be central to syt’s function in triggering Ca^2+^ induced exocytosis (van den Bogaart *et al*., 2012).

In this paper we developed a mathematical model in which the dual binding of the C2B domain to Ca^2+^ and PI(4,5)P_2_ is promoting fusion. The model, which is based on the measured affinities and allostericity of Ca^2+^ and PI(4,5)P_2_ binding, describes stochastic binding/unbinding reactions at the level of individual syts and stochastic SV fusion events. The model predicts that each C2B domain engaging in dual Ca^2+^/PI(4,5)P_2_ binding lowers the energy barrier for fusion by 4.85 k_B_T. Three vesicular syts simultaneously engage their C2B domains in dual Ca^2+^/PI(4,5)P_2_ binding for fast NT release and this simultaneous engagement of multiple syts crucially relies on the positive allosteric coupling between Ca^2+^ and PI(4,5)P_2_ binding. We explored consequences of putative mutants affecting Ca^2+^/PI(4,5)P_2_ binding and/or the allosteric coupling between the binding sites of both species and suggest that changes of the allostericity contribute to dominant negative effects. Moreover, dynamic changes of PI(4,5)P_2_ accessibility for the syts (e.g. induced by SV movement to the plasma membrane) are predicted to dramatically impact synaptic responses. We conclude that allosterically stabilized Ca^2+^/PI(4,5)P_2_ dual binding to the C2B domains forms the molecular basis for synaptotagmins to exert their cooperative control of neurotransmitter release.

## Results

### An experiment-based model of the triggering mechanism for SV fusion based on molecular interactions between syt, Ca^2+^, and PI(4,5)P_2_

To develop an experiment-based model of NT release based on molecular properties of syt, we first described the reaction scheme of a single C2B domain. The C2B domain binds PI(4,5)P_2_ and two Ca^2+^ ions (Fernández-Chacón et al., 2002; Honigmann *et al*., 2013; Mackler *et al*., 2002; van den Bogaart *et al*., 2012; Xue *et al*., 2008). Therefore, in our model the C2B domain can be in four different states (Figure 1A): (1) an “empty” state, (2) a PI(4,5)P_2_-bound state, (3) a state with two Ca^2+^ ions bound, and (4) a “dual-bound” state in which the C2B simultaneously engages Ca^2+^/PI(4,5)P_2_ binding. The binding affinities of syt1 C2B for Ca^2+^ and PI(4,5)P_2_ were set to those measured *in vitro* (K_D,2Ca2+_ and K_D,PIP2_)(van den Bogaart *et al*., 2012). Binding of PI(4,5)P_2_ to the C2B domain was shown to increase the domain’s affinity for Ca^2+^ and vice versa, indicating a positive allosteric coupling between the two binding sites (van den Bogaart *et al*., 2012). We therefore implemented a positive allosteric stabilization of the dual-bound state in the model (illustrated by the red shaded areas of the C2B domain in Figure 1A) by introducing the allosteric factor (A= 0.00022, see <u>Methods</u> (van den Bogaart *et al*., 2012)) which slows down the Ca^2+^ and PI(4,5)P_2_ dissociation from the dual-bound state.

**Figure 1:**
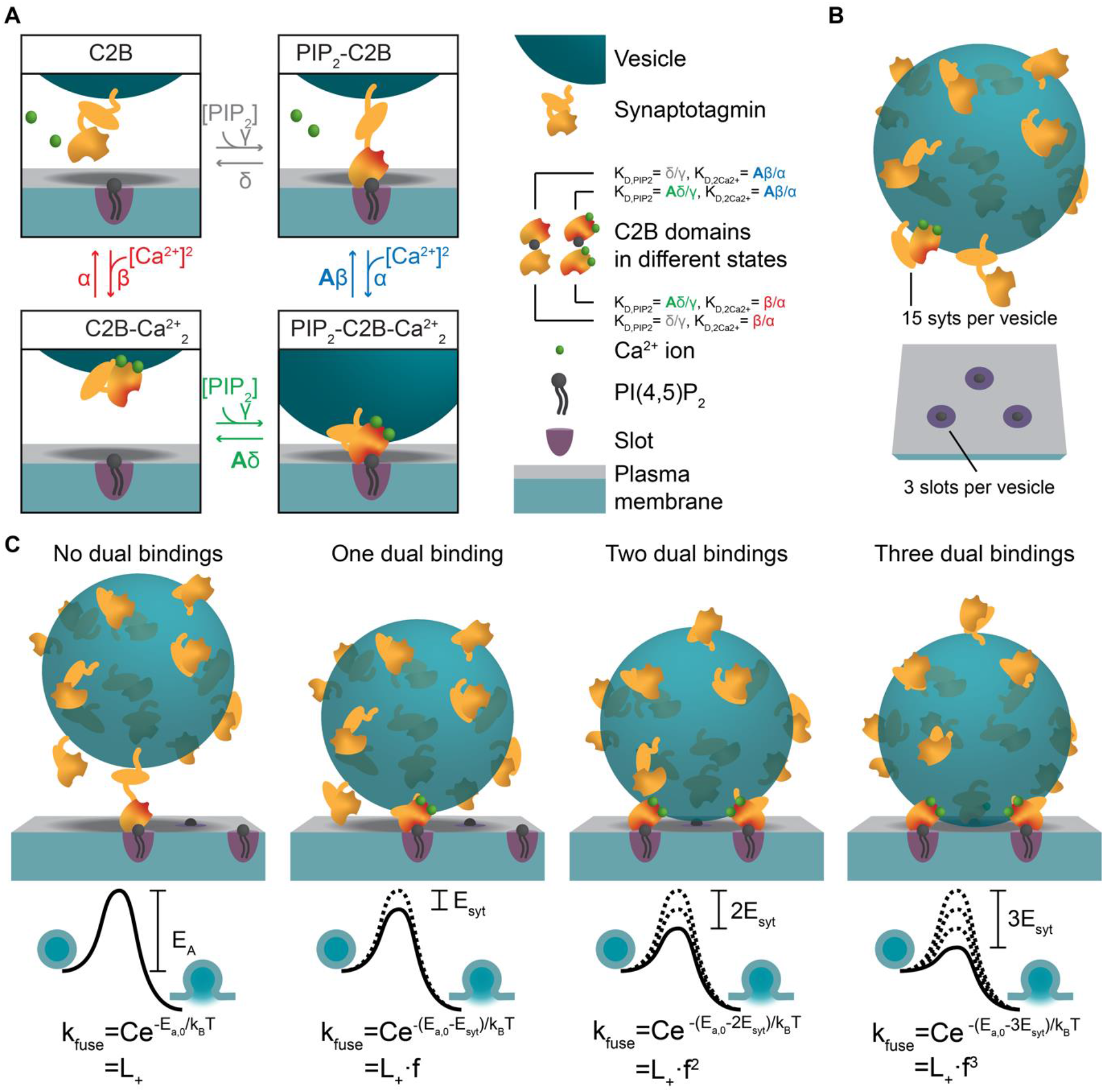
A molecular model of NT release triggered by Ca^2+^ and PI(4,5)P_2_ binding to the syt1 C2B domain. **A)** The reaction scheme of a single syt C2B domain. Each syt can be in one of four binding states: Nothing bound (top left), PI(4,5)P_2_ bound (top right), two Ca^2+^ ions bound (bottom left), and PI(4,5)P_2_ and two Ca^2+^ ions bound (bottom right). Simultaneous binding of Ca^2+^ and PI(4,5)P_2_ to the syt C2B domain is referred to as dual binding. The factor A<1 on the dissociation rates (β and δ) from the dual-bound state represents the positive allosteric effect of simultaneous PI(4,5)P_2_ and Ca^2+^ binding and leads to stabilization of the dual-bound state. The ratio between dissociation rate and association rate constants (β/α and δ/γ) is equal to the respective dissociation constants of syt1 determined *in vitro* (K_D,2Ca2+_ =221^2^ µM^2^ and K_D,PIP2_ =20 µM, (van den Bogaart *et al*., 2012)). An alternative reaction scheme where Ca^2+^ binding leads to association of the C2B domain with the plasma membrane is shown in Figure 1 – figure supplement 1. Our model is not influenced by the assumptions on whether Ca^2+^ binding to syt leads to plasma membrane or vesicle association. **B)** The stoichiometry at the SV fusion site. We assume 15 syts per SV (Takamori *et al*., 2006), and that the association of the syt C2B domain to PI(4,5)P_2_ is limited to a finite number of slots (here illustrated for *M*_*slots*_=3). **C)** The effect of formation of multiple dual bindings on the energy barrier for SV fusion and the SV fusion rate. We assume that each dual-binding C2B domain lowers the energy barrier for fusion by the same amount (E_syt_, illustrated in middle row), thereby increasing the fusion rate (k_fuse_) with a factor *f* for each dual binding (equation in bottom row). The fusion rate for an SV with no dual bindings formed is set to L_+_. The model is a Markov model, which can be summarized in a state diagram describing the reactions of the syt-harboring SV (Figure 1 – figure supplement 2).

We next extended the model to the level of the complete SV. On average, SVs contain 15 copies of syt1 (Figure 1B) (Takamori et al., 2006), which may work together to regulate SV fusion. Spontaneous release occurs at low rates (with a rate constant ‘L_+_’), reflected by a high initial energy barrier for SV fusion (Figure 1C). Because syt’s stimulation of SV fusion likely relies on the simultaneous binding of both Ca^2+^ and PI(4,5)P_2_ (Kedar *et al*., 2015; Li *et al*., 2006; Mackler *et al*., 2002; Mackler and Reist, 2001; Wang *et al*., 2016; Wu *et al*., 2021a; Wu *et al*., 2021b), we assumed that each dual-bound C2B domain promotes exocytosis by lowering this barrier. How this might be achieved exactly is unknown, but could involve bridging plasma and SV membranes (Figure 1A), changing the curvature of the plasma membrane (Figure 1 – Figure supplement 1), changing the local electrostatic environment, or directly or indirectly promoting SNARE complex assembly (Bhalla et al., 2006; Martens et al., 2007; Ruiter et al., 2019; Schupp *et al*., 2016; Tang et al., 2006; van den Bogaart et al., 2011b; Zhou *et al*., 2015; Zhou *et al*., 2017). We assumed that multiple syts may progressively lower the energy barrier by the successive engagement of their C2B domains in dual Ca^2+^/PI(4,5)P_2_ binding, and investigated the simplest scenario, in which each dual-bound C2B domain contributed the same amount of energy (*E*_*syt*_) thereby increasing the fusion rate by the same factor (*f*) (Figure 1C)(Schotten et al., 2015). The number of syts that can simultaneously promote fusion may be limited by their access to PI(4,5)P_2_ in the plasma membrane. This could be due to limited space beneath the SV or limited molecular access to bind PI(4,5)P_2_ clusters and/or SNAREs (de Wit *et al*., 2009; Mohrmann *et al*., 2013; Rickman and Davletov, 2003). We therefore created a model that stochastically describes the binding status of individual SVs (Figure 1 – Figure supplement 2) with a limited number of PI(4,5)P_2_ association possibilities (‘slots’) and investigated how this number affects physiological responses.

### At least three PI(4,5)P_2_ binding slots are required to reproduce release kinetics from the calyx of Held synapse

A hallmark of the NT release reaction is its large dynamic range in response to Ca^2+^ stimuli as impressively demonstrated by experimental data from the calyx of Held synapse where release latencies (defined as the time of the fifth SV fusion after the stimulus) and exocytosis rates have been measured for a broad range of Ca^2+^ concentrations (using Ca^2+^ uncaging)(Bollmann *et al*., 2000; Kochubey and Schneggenburger, 2011; Lou *et al*., 2005; Schneggenburger and Neher, 2000; Sun et al., 2007). At this well-established model synapse, fast NT release is controlled by syt2, which is functionally redundant with syt1 in neurons (Kochubey *et al*., 2016; Xu et al., 2007).

We evaluated whether our model could reproduce this Ca^2+^ dependence by simulating release latencies and peak release rates in response to step-like Ca^2+^ stimuli. The ability to reproduce the experimental data depended on the number of ‘slots’ for syt PI(4,5)P_2_ binding (Figure 2A). We first fitted the free parameters in our model by optimizing the agreement (i.e. by reducing a pre-defined cost function, see <u>Methods</u>) between model predictions and release rates and latencies determined experimentally by Kochubey and Schneggenburger (Figure 2A)(Kochubey and Schneggenburger, 2011). During this fitting process, we took the entire distribution of the experimentally obtained release latencies into account by using the likelihood function (see <u>Methods</u>). This was not feasible for the experimental peak release rates (since accurate computation of the maximum rate of stochastic events is not feasible), which were therefore compared to the closed form solution of the model (see <u>Methods</u>). Because the affinities for Ca^2+^ and PI(4,5)P_2_, and the allosteric coupling between both species (K_D,2Ca2+_, K_D,PIP2_ and *A*, Figure 1A, Figure 1 – figure supplement 2) as well as the RRP size (including its variance) were taken from literature (Table 1) (Figure 2 – figure supplement 1) (van den Bogaart *et al*., 2012; Wölfel and Schneggenburger, 2003), we only had to estimate five unknown (“free”) parameters: (1) the binding rate constant of the two Ca^2+^ ions (*α*), (2) the binding rate constant of PI(4,5)P_2_ (*γ*), (3) the PI(4,5)P_2_ concentration ([PI(4,5)P_2_]), (4) the reduction of the energy barrier for fusion induced by Ca^2+^/PI(4,5)P_2_ dual binding of one C2B domain (*E*_*syt*_) and (5) (like in previous models) a fixed delay (*d*) between time of uncaging and response onset (see (Kochubey and Schneggenburger, 2011; Schneggenburger and Neher, 2000)). Having obtained the best fit parameters, peak release rates and release latencies (Figure 2A) were stochastically simulated based on the closed-form solution of the Markov model (see <u>Methods</u>).

**Figure 2:**
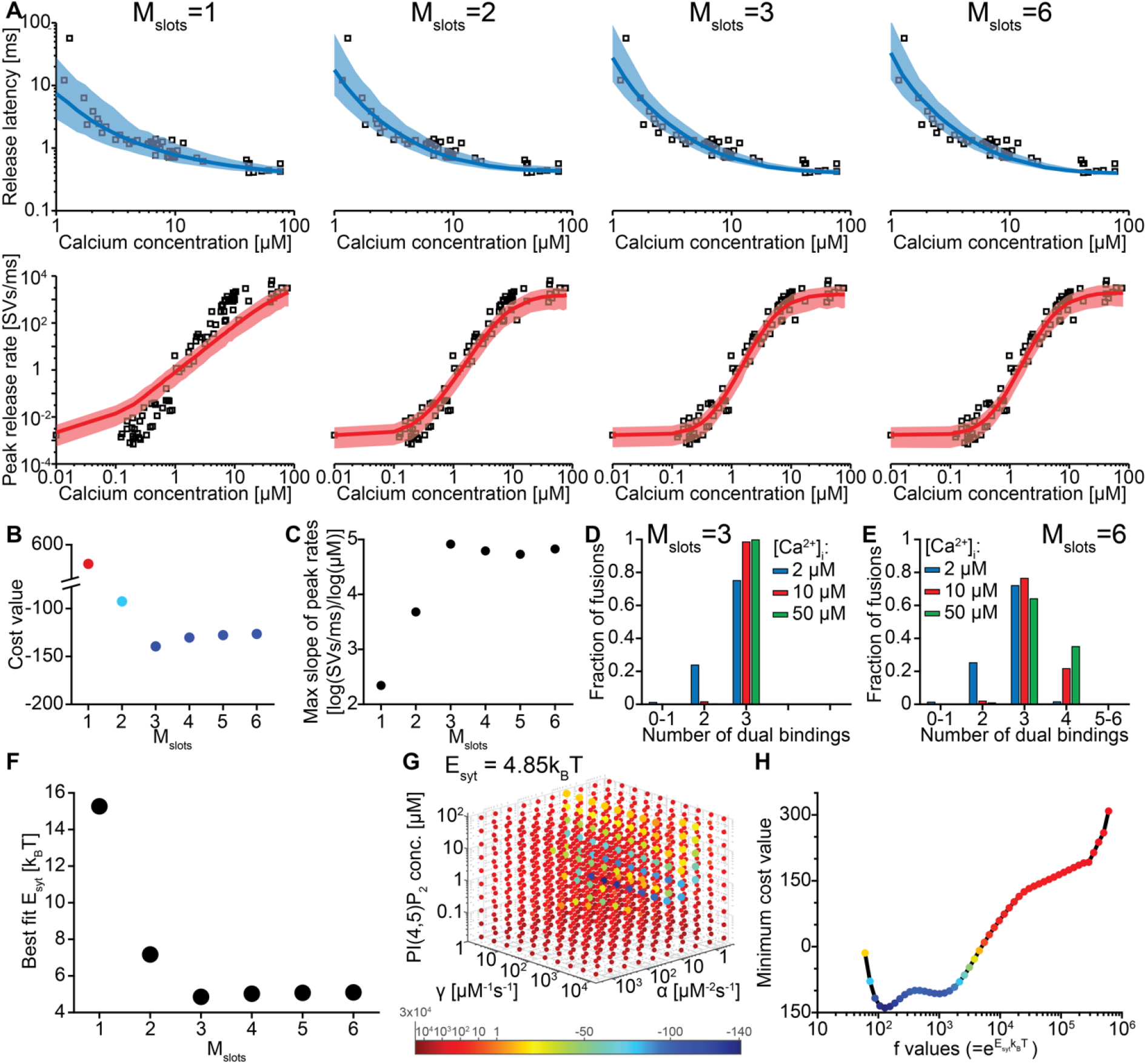
The model reproduces the Ca^2+^ dependency of SV fusion when at least three syts can simultaneously bind PI(4,5)P_2_. **A)** Best fit results for different choices of *M*_*slots*_. The top panels show best fit model prediction of the release latencies (time to fifth SV fusion), and the bottom panels show the predicted peak release rates at varying Ca^2+^ concentrations. The black points are experimental data (individual measurements replotted from Kochubey and Schneggenburger (2011)). Solid lines represent the median release latencies and mean peak release rates predicted by the model from 1000 repetitions per simulated [Ca^2+^]_i_. The shaded areas indicate the 95% prediction interval of the model. The models with *M*_*slots*_<3 failed to reproduce data, whereas models with *M*_*slots*_≥3 agreed with data. Optimization and simulation were performed using a variable RRP size (Figure 2 – figure supplement 1). **B)** Minimum cost value as a function of *M*_*slots*_. With *M*_*slots*_=3 the minimum cost value was obtained, indicating the best correspondence to experimental release latencies and peak release rates. The point colors correspond to the color scale in panel G. **C)** Maximal slope of logarithm of simulated peak release rate vs logarithm of [Ca^2+^]_i_. For *M*_*slots*_<3 the model failed to reproduce the Ca^2+^ dependency of release rates. **D-E)** The number of dual bindings at the time point of fusion for *M*_*slots*_=3 (D) and *M*_*slots*_=6 (E) determined from simulations of 10^4^ SVs using three different [Ca^2+^]_i_. Most fusions took place after forming 3 or 4 dual bindings, even when allowing more dual bindings to form. A larger set of Ca^2+^ stimuli was also explored (Figure 2 – figure supplement 2 and Figure 2 - figure supplement 3). Forcing a model with M_slots_=6 to fuse after 5/6 dual bindings were formed could not describe the experimental data and showed a too steep Ca^2+^ dependency of release (Figure 2 – figure supplement 4). **F)** The change in the energy barrier induced by dual binding formation (E_syt_) as a function of *M*_*slots*_. E_syt_ was computed from the fitted *f* values and was approximately constant for *M*_*slots*_≥3. **G)** Exploring cost values in the parameter space for a model with *M*_*slots*_ =3. With *f* fixed at the best fit value (*f*=128), we determined the cost value of all combinations of 30 choices of the three free parameters, α, γ and [PI(4,5)P_2_]. As the added delay only leads to a vertical shift in the release latencies plot (see Figure 2 – figure supplement 5), this parameter was optimized for each choice of the other free parameters to minimize the costs. The plot shows a subset of the parameter combinations, and the colors indicate the cost value at each point. The color scale is linear below 1 and logarithmic above 1, and points with a cost value >1 are smaller for better visibility. The darkest blue colored ball represents the overall minimum cost value in this parameter search and agrees with the best fit obtained. The effect of varying each of the free parameters on release latencies and peak release rates can be seen in Figure 2 – figure supplement 5. **H)** Minimum cost value as a function of *f* for a model with *M*_*slots*_ =3. For each choice of *f* the model was fit to experimental data. This parameter exploration found the same minimum in the parameter space as found by fitting all free parameters. Simulation scripts can be found in Source code 1. Results depicted in Figure 2 and its figure supplements can be found in Figure 2 - source data 1.

**Table 1.** Fixed parameters in the model.

| Parameter | Description | Value | Reference |
| --- | --- | --- | --- |
| $n_{syts}$ | Number of syts per SV | 15 | (Takamori *et al.*, 2006) |
| $M_{slots}$ | Number of binding slots for syt to PI(4,5)P<sub>2</sub> (see Figure 2) | Varied from 1-6 | This paper |
| $n_{ves}$ | Number of RRP vesicles | Mean: 4000, sd: 2000, gamma distribution | (Wölfel and Schneggenburger, 2003) |
| $[Ca^{2+}]_0$ | Resting $[Ca^{2+}]_i$ | 0.05 μM | (Helmchen *et al.*, 1997) |
| $K_{D,2Ca^{2+}}$ | Dissociation constant of C2B for two $Ca^{2+}$ ions | $221^2 \mu M^2$ | (van den Bogaart *et al.*, 2012) |
| $K_{D,PIP2}$ | Dissociation constant of C2B for PI(4,5)P<sub>2</sub> | 20 μM | (van den Bogaart *et al.*, 2012) |
| $A$ | Allosteric factor see Methods for calculation | 0.00022 | (van den Bogaart *et al.*, 2012) |
| $\beta$ | $Ca^{2+}$ unbinding rate constant | $K_{D,2Ca^{2+}} \cdot \alpha$ | Computed using best fit $\alpha$ (see Table 2) |
| $\delta$ | PI(4,5)P<sub>2</sub> unbinding rate constant of C2B | $K_{D,PIP2} \cdot \gamma$ | Computed using best fit $\alpha$ (see Table 2) |
| $L_+$ | Basal fusion rate | $4.23 \cdot 10^{-4} s^{-1}$ | Computed using data from (Kochubey and Schneggenburger, 2011) |

We systematically varied the number of slots (*M*_*slots*_) from one to six, optimized the free parameters for each of these choices, and compared the best fit solutions. With a single slot (*M*_*slots*_ =1), the model accounted for experimentally observed release latencies, but failed to reproduce the steep relationship between [Ca^2+^]_i_ and peak release rates (Figure 2A). Adding more slots strongly improved the agreement with experimental data. The best agreement was found with three slots (*M*_*slots*_=3) and the agreement slightly decreased with more slots (*M*_*slots*_ >3) (Figure 2B). However, all models with at least three slots (*M*_*slots*_≥3) captured the steep dependency of peak release rates on [Ca^2+^]_i_, with a maximum slope of 4-5 on a double-logarithmic plot (Figure 2C) (Schneggenburger and Neher, 2000).

Our model made it possible to inspect the fate of each individual fusing SVs, including the number of synaptotagmins dually binding Ca^2+^ and PI(4,5)P_2_ just before fusion. Remarkably, even in models with more than three slots (*M*_*slots*_>3), fast NT release (for moderate to high [Ca^2+^]_i_) was predicted to primarily engage three dual-bound syt C2B domains (while fewer dual-bound syts mediated fusion for lower [Ca^2+^]_i_)(Figure 2D-E, Figure 2 - figure supplement 2, Figure 2 - figure supplement 3). Correspondingly, the estimated reduction of the energy barrier for fusion by each Ca^2+^/PI(4,5)P_2_ dual-binding C2B domain was similar (E_syt_≈5 k_B_T) for all versions of the model with at least three slots (M_slots_≥3) (Figure 2F, see Table 2 for other best fit model parameters). Manually forcing a lower contribution to the energy barrier for fusion in a model with six slots and refitting the remaining parameters, which would be required to involve more dual bindings in fusion, resulted in a too steep dependence of the peak release rate vs. [Ca^2+^]_I_ (figure 2 – figure supplement 4). Thus, our model predicts that three syts actively reduce the energy barrier for fast SV fusion. Consequently, we used the model with three slots (*M*_*slots*_=3) for further simulations.

**Table 2.**
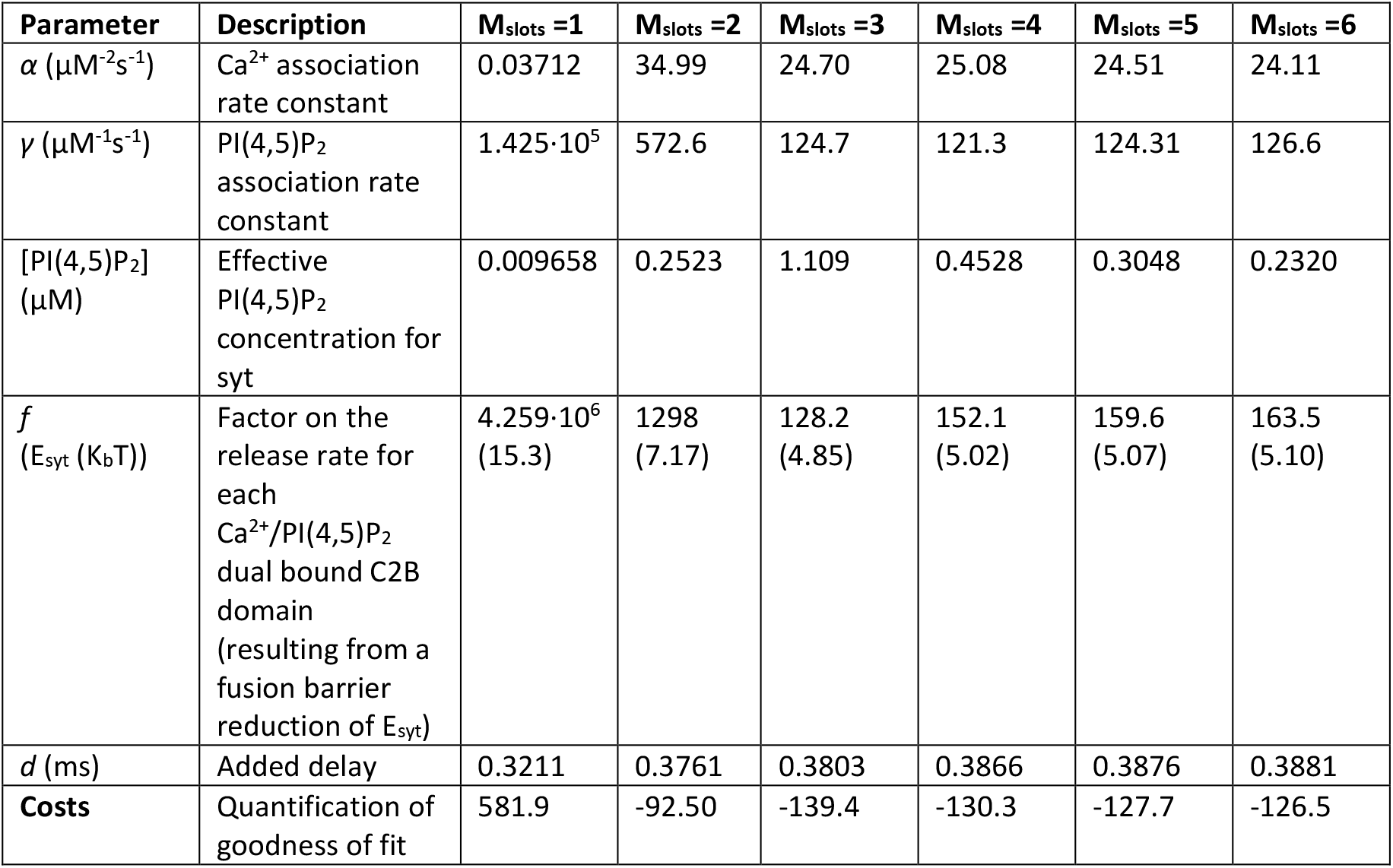
Best fit model parameters and corresponding costs with different number of slots.

| Parameter | Description | M<sub>slots =1</sub> | M<sub>slots =2</sub> | M<sub>slots =3</sub> | M<sub>slots =4</sub> | M<sub>slots =5</sub> | M<sub>slots =6</sub> |
| --- | --- | --- | --- | --- | --- | --- | --- |
| $\alpha$ ($\mu\text{M}^{-2}\text{s}^{-1}$) | Ca<sup>2+</sup> association rate constant | 0.03712 | 34.99 | 24.70 | 25.08 | 24.51 | 24.11 |
| $\gamma$ ($\mu\text{M}^{-1}\text{s}^{-1}$) | PI(4,5)P<sub>2</sub> association rate constant | 1.425·10<sup>5</sup> | 572.6 | 124.7 | 121.3 | 124.31 | 126.6 |
| [PI(4,5)P<sub>2</sub>] ($\mu\text{M}$) | Effective PI(4,5)P<sub>2</sub> concentration for syt | 0.009658 | 0.2523 | 1.109 | 0.4528 | 0.3048 | 0.2320 |
| $f$ (E<sub>syt</sub> (K<sub>b</sub>T)) | Factor on the release rate for each Ca<sup>2+</sup>/PI(4,5)P<sub>2</sub> dual bound C2B domain (resulting from a fusion barrier reduction of E<sub>syt</sub>) | 4.259·10<sup>6</sup> (15.3) | 1298 (7.17) | 128.2 (4.85) | 152.1 (5.02) | 159.6 (5.07) | 163.5 (5.10) |
| $d$ (ms) | Added delay | 0.3211 | 0.3761 | 0.3803 | 0.3866 | 0.3876 | 0.3881 |
| **Costs** | Quantification of goodness of fit | 581.9 | -92.50 | -139.4 | -130.3 | -127.7 | -126.5 |

The best fit parameters for three slots revealed rapid association rate constants for Ca^2+^ and PI(4,5)P_2_ to the C2B domain and PI(4,5)P_2_ levels corresponding to a concentration of ∼1 µM in an in vitro setting (Table 2). Predicted responses obtained using the best-fit parameters were sensitive to changes of either of these parameters (Figure 2 – figure supplement 5). For instance, higher levels of PI(4,5)P_2_ decreased the release latencies and increased the rate of fusion, and changing the Ca^2+^ association rate constant (*α*) affected the release latencies much more than changing the PI(4,5)P_2_ association rate constant (*γ*). We verified that these parameters represent unique solutions by systematically exploring the parameter space with *f* (which relates to the lowering of the fusion barrier for each syt dual-binding Ca^2+^/PI(4,5)P_2_) fixed to the best fit value (*f*=128), which revealed a clear minimum at the best fit parameters (Figure 2G, darkest ball). We furthermore confirmed that this *f* value was optimal by systematically varying *f* and fitting all other parameters (Figure 2H).

### The number of syt proteins pre-associated to PI(4,5)P_2_ at rest influences the SV’s Ca^2+^ responsiveness

The steady state concentration of PI(4,5)P_2_ determines the probability of syts associating to PI(4,5)P_2_ at rest. With the best fit parameters, our model predicts that at rest ([Ca^2+^]_i_=50 nM) most SVs associate to PI(4,5)P_2_ by engaging one (∼42%), two (∼33%) or three (∼8%) syts (Figure 3A, see Figure 3 – figure supplement 1 for behavior in the model with *M*_*slots*_=6). With a step-like Ca^2+^ stimulus to 50 µM, SVs with two or three pre-associated syts mediated most of the fastest (<0.5 ms) SV fusions (Figure 3B). Consequently, changing the steady state PI(4,5)P_2_ concentration (which changes the number of pre-associated syts/SVs) largely impacted the release latencies (defined as the timing of the fifth SV that fuses (Kochubey and Schneggenburger, 2011) Figure 2 – figure supplement 5). Due to the allosteric coupling, the Ca^2+^ affinities of syts prebound to PI(4,5)P_2_ are increased and SVs with more PI(4,5)P_2_ interactions are more responsive to the Ca^2+^ stimulus (Figure 3C). Thus, at the single SV level, the number of pre-associated syts to PI(4,5)P_2_ at rest plays a role in very fast (submillisecond) SV release and causes heterogeneity in release probability among RRP SVs (Wolfel et al., 2007).

**Figure 3:**
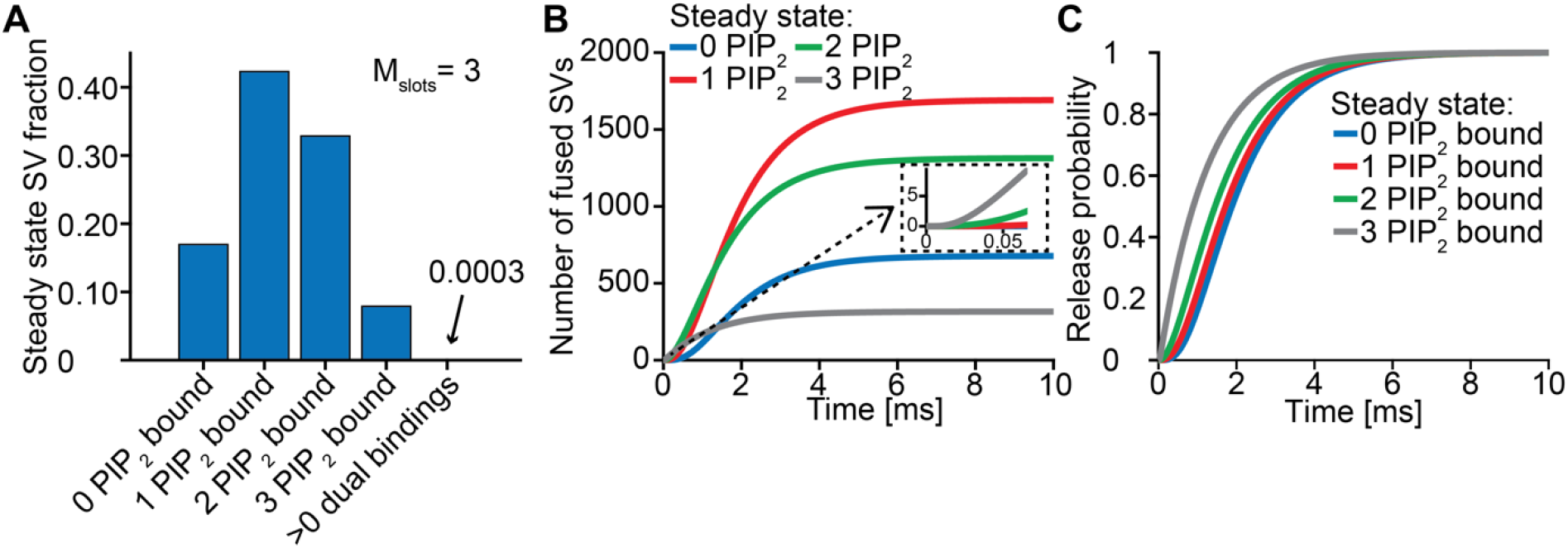
Syts binding to PI(4,5)P_2_ prior to Ca^2+^ stimulus underlies very fast SV fusion. **A)** PI(4,5)P_2_ binding status of SVs at steady state. At resting [Ca^2+^]_i_ of 50 nM, more than 40 % of SVs have bound a single PI(4,5)P_2_ molecule, more than 30 % have bound two PI(4,5)P_2_, while less than 10 % have bound three PI(4,5)P_2_. Close to no SVs form dual bindings at steady state. **B)** Cumulative fusion of SVs after 50 µM step Ca^2+^ at t=0, grouped according to their initial PI(4,5)P_2_ binding state. During the first ∼0.5 ms, release is dominated by SVs having two or three syts bound to PI(4,5)P_2_ prior to the stimulus. The insert shows that the SVs having prebound three PI(4,5)P_2_ constitute the majority of the first five SVs that fuse in response to the Ca^2+^ step and therefore have largely impacted the release latency. **C)** Cumulative release probability of SVs over time after 50 µM step Ca^2+^ at t=0, grouped according to initial PI(4,5)P_2_ binding state. The dominance of SVs having pre-bound to PI(4,5)P_2_ with two or three syts in panel B is explained by their high release probability compared to SVs with no or only one PI(4,5)P_2_ bound. Figure 3 - figure supplement 1 shows the same analysis for a model with M_slots_= 6. Simulation scripts can be found in Source code 1. Depicted simulation results can be found in Figure 3 - source data 1.

### Allosteric stabilization of Ca^2+^/PI(4,5)P_2_ dual binding is necessary to synchronize multiple C2B domains for fast SV fusion

An important feature of our model is the inclusion of a positive allosteric interaction between Ca^2+^ and PI(4,5)P_2_ binding to the C2B domain which we based on increased affinities measured *in vitro* (van den Bogaart *et al*., 2012). To explore the physiological relevance of this allostericity, we investigated how individual SVs engaged their syt C2B domains in Ca^2+^/PI(4,5)P_2_ dual binding in response to a stepwise increase of [Ca^2+^]_i_ to 50 µM (Figure 4A) with or without this allosteric coupling. We did this by following the fate of four randomly chosen RRP SVs in stochastic simulations (Figure 4B). Under normal conditions (with allostericity) syt C2B domains quickly associated both Ca^2+^ and PI(4,5)P_2_ and their respective allosteric stabilization slowed the dissociation of either species resulting in a lifetime of their dual binding of ∼1.3 ms on average. This enabled the successive engagement of three dual-bound C2B domains at all four investigated SVs which induced their fusion within 4 ms (the average waiting time for fusion with three dual-bound syts was ∼1.1 ms)(Figure 4B, fusion indicated by circles). By inspecting the average behavior of the entire RRP of SVs it became clear that the overall release rate closely followed the population of SVs engaging three C2B domains in dual Ca^2+^/PI(4,5)P_2_ binding, illustrating the importance of engaging three syts for fast SV fusion (Figure 4C). We also simulated the postsynaptic response produced by this NT release by convolving the SV release rate with a typical postsynaptic response to the fusion of a single SV (see <u>Methods</u>), which revealed synchronous and large Excitatory Post Synaptic Currents (EPSCs)(Figure 4D).

**Figure 4:**
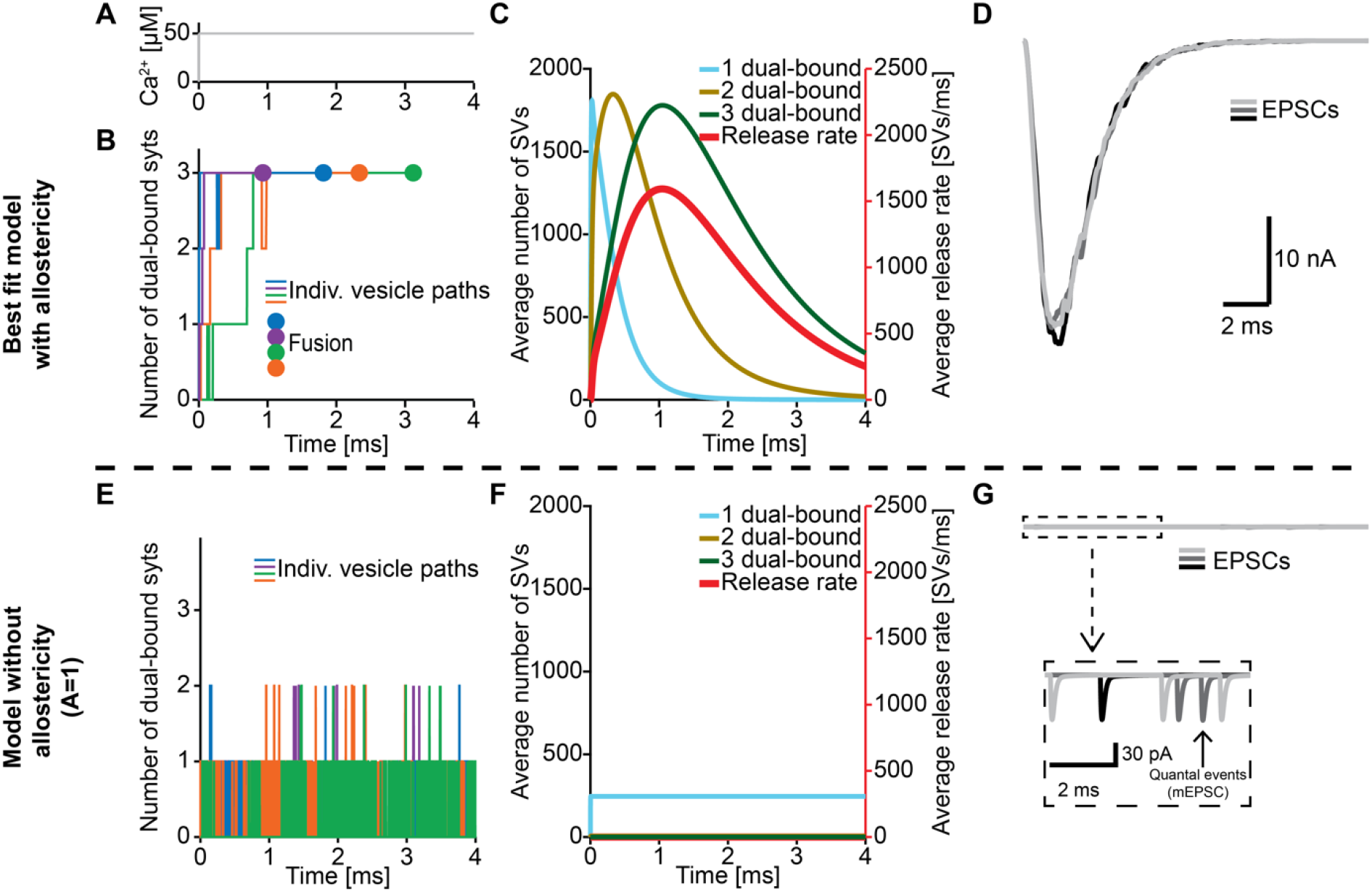
The positive allostericity between Ca^2+^ and PI(4,5)P_2_ allows multiple syt C2B domains to engage in in Ca^2+^/PI(4,5)P_2_ dual binding. **A)** Ca^2+^ signal used in simulations ([Ca^2+^]_i_=50 µM). This constant Ca^2+^ concentration was used for all simulations depicted in this figure. **B)** The path towards fusion for four example SVs using stochastic simulations of the best fit model (with allostericity). The differently colored graphs show the number C2B domains engaging in Ca^2+^/PI(4,5)P_2_ dual binding for the four example SVs. The large dots indicate SV fusion. **C)** Average number of SVs having one (blue), two (olive) and three (green) C2B domains engaging in Ca^2+^/PI(4,5)P_2_ dual binding and the fusion rate (red) over time in simulations including the entire RRP. In the best fit model, the number of SVs with three syts dual-binding Ca^2+^/PI(4,5)P_2_ peaks approximately at the same time as the fusion rate. The decrease in number of SVs with one or two C2B domains dual binding Ca^2+^/PI(4,5)P_2_ reflects formation of additional dual bindings. The decrease in total number of SVs is caused by fusion of RRP vesicles. **D)** Excitatory Postsynaptic Currents (EPSCs) from three stochastic simulations with a fixed RRP size of 4000 SVs. The model predicts synchronous EPSCs with a small variation caused by the stochasticity of the molecular reactions. **E)** The path towards fusion for four example SVs (similar to panel A) in the model without allostericity in stochastic simulation. All parameters other than the allosteric factor (*A*) are the same as in the best fit model. Without stabilization, the syts quickly engage *and* disengage in Ca^2+^/PI(4,5)P_2_ dual binding and rarely more than one syt engages in dual binding. Formation and dissociation of Individual dual-bound syts is too fast to distinguish on the time scale depicted here. No fusions occurred in the depicted simulations. **F)** Average number of SVs having one (blue), two (olive) and three (green) syts engaging in Ca^2+^/PI(4,5)P_2_ dual binding and the fusion rate (red) over time in the model without allostericity. Almost no SVs form more than one syt dual binding Ca^2+^/PI(4,5)P_2_, which results in a very low fusion rate. **G)** EPSCs from three stochastic simulations and with a fixed RRP size of 4000 SVs. A model lacking allostericity only shows sporadic, individual release events. The insert shows a zoom-in of the first 6 ms of simulation and makes single SV fusion events giving rise to quantal, ‘miniature’ mEPSCs visible. Fitting the model without the allosteric effect to the experimental data was unsuccessful (Figure 4 – figure supplement 1). Simulation scripts can be found in Source code 1. Results from fitting the model without allosteric effect can be found in Figure 4 - source data 1.

We then explored what would happen without the allosteric stabilization of Ca^2+^/PI(4,5)P_2_ dual binding (by setting A=1; Figure 4E). In this case, the C2B domains still quickly associated Ca^2+^ and PI(4,5)P_2_, but without the allosteric slowing of Ca^2+^/PI(4,5)P_2_ dissociation the lifetime of dual-bound C2B domains was dramatically reduced (to an average of ∼0.0003 ms). This made it very improbable to engage multiple C2B domains in dual Ca^2+^/PI(4,5)P_2_ binding (Figure 4E). In turn, without the simultaneous engagement of multiple syts dual-binding Ca^2+^/PI(4,5)P_2_, NT release became very unlikely. In fact, none of the randomly chosen four RRP SVs fused within 4 ms (Figure 4E). Inspection of the average behavior of the entire RRP revealed that only few SVs engaged more than one syt C2B domain in dual Ca^2+^/PI(4,5)P_2_ binding, resulting in a very low fusion rate (Figure 4F). Correspondingly, postsynaptic EPSCs were severely disrupted, and most release events were ill-synchronized single SV fusion events (Figure 4G). It was furthermore not possible to fit a model without the allosteric stabilization to the experimental dataset (Figure 4 – figure supplement 1). Thus, the positive allosteric coupling between Ca^2+^ and PI(4,5)P_2_ is fundamental for the syts to simultaneously and persistently engage multiple C2B domains per SV in Ca^2+^/PI(4,5)P_2_ dual binding.

### Many syt per SV speed up exocytosis by increasing the probability of Ca^2+^/PI(4,5)P_2_ dual binding

Our model strongly suggests that no more than three syts simultaneously binding Ca^2+^ and PI(4,5)P_2_ are required to promote fast SV fusion (Figure 2). Yet, a total of 15 copies are expressed per SV on average (Takamori *et al*., 2006), which raises the question why SVs carry such excess and whether and how the additional syt copies contribute to the characteristics of Ca^2+^-induced synaptic transmission. To investigate this, we simulated Ca^2+^ uncaging experiments with reduced numbers of syts per SV while keeping all other parameters in the model constant. With fewer syts, release latencies increased and peak release rates reduced. Defects were particularly prominent for reductions to less than three copies per SV (Figure 5A). Further exploration indicated that this was because it took SVs longer to simultaneously engage three C2B domains in dual Ca^2+^/PI(4,5)P_2_ binding and that fewer SVs reached this state (Figure 5B).

**Figure 5:**
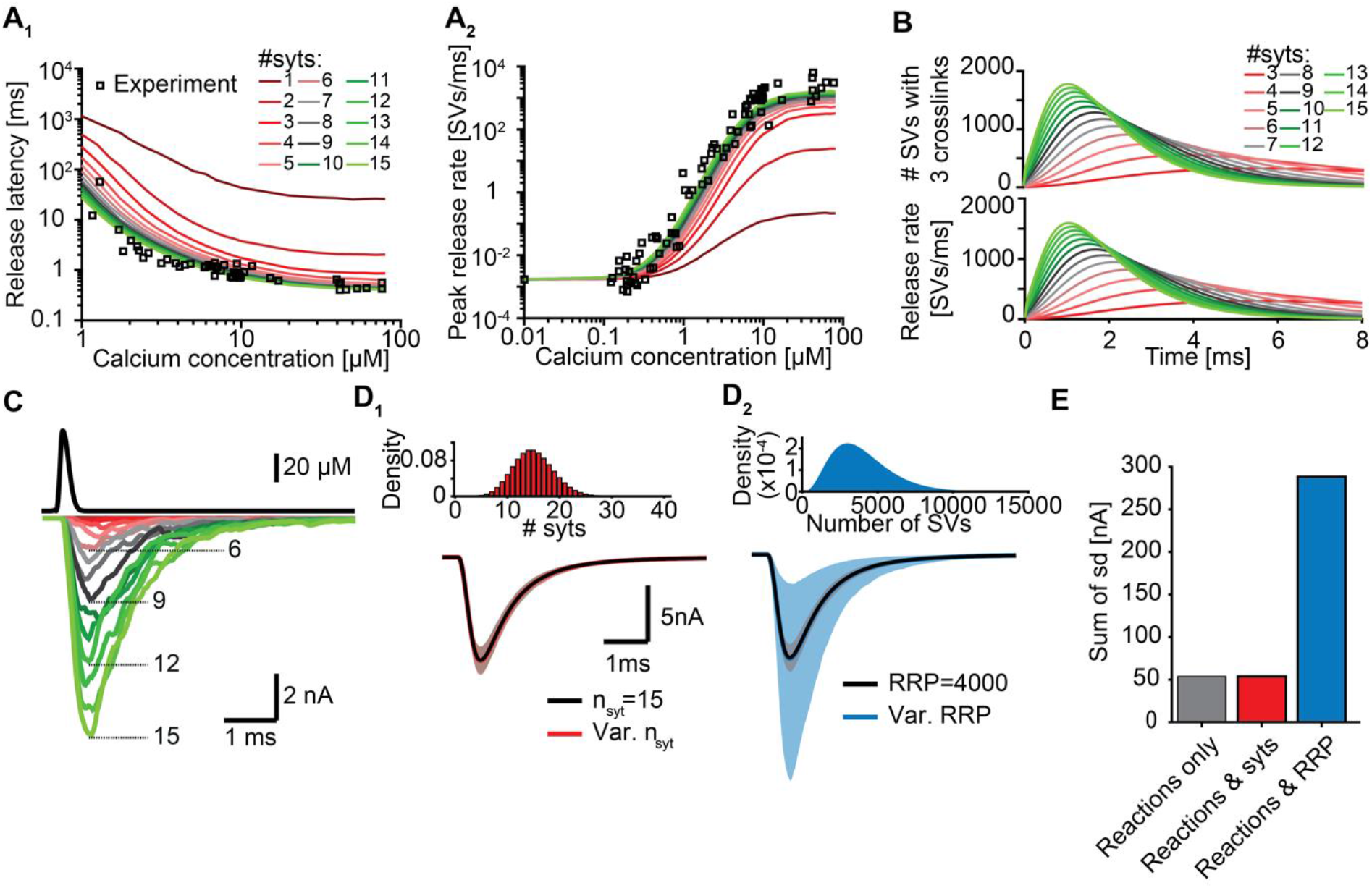
Simulations with reduced syt expression predict a reduction in SV fusion. **A)** Model predictions of median release latencies (A_1_) and mean peak release rates (A_2_) as a function of [Ca^2+^]_i_ for different numbers of syts per SV. All simulations were performed with 1000 repetitions using the best fit parameters obtained by fitting with *n*_*syts*_ = 15. Experimental data points are replotted from (Kochubey and Schneggenburger, 2011). **B**) The average number of SVs with three dual bindings formed (top) and release rate (bottom) as a function of time for 3-15 syt copies per SV from simulations with a Ca^2+^ flash of 50 µM. **C)** Predicted AP-evoked responses (bottom) simulated using a realistic Ca^2+^ transient (top) (Wang et al., 2008) for different numbers of syts per SV. The AP-evoked response shown at the bottom are representative single stochastic simulations with an amplitude closest to the mean amplitude of 200 repetitions. Increasing [PI(4,5)P_2_] for each choice of *n*_*syt*_ ≥ 3 could fully rescue release latencies, release rates, and evoked responses (Figure 5 – figure supplement 2). **D)** Variability in simulated AP-evoked responses for a model with a variable number of syts and an RRP size of 4000 (D_1_, red, bottom) and a model with a variable RRP size and 15 syts per SV (D_2_, blue, bottom) compared to the variance induced by the stochasticity of the reactions only (with fixed number of SVs and syts, grey). Solid lines depict mean traces and the shaded area indicates the 95% prediction interval. Simulations with 1000 repetitions. Top panels show the probability density distributions of the number of syts (Poisson distribution, lambda =15) and of the number of SVs (gamma distribution, mean of 4000 SVs, standard deviation of 2000 SVs, outcome rounded to nearest integer). **E)** Quantification of the variance in the traces introduced by the stochasticity of the model reactions (grey), by the stochasticity of model reactions and variable syt number (red), and by the stochasticity of model reactions and variable RRP size (blue) by computing the sum of the standard deviation (sd) determined over the entire trace (0-6 ms, 300 data points). Simulation of the individual syts using the Gillespie algorithm agreed with simulations using the analytical solution of the Markov model (Figure 5 – figure supplement 1 and Methods). Simulation scripts can be found in Source code 1. Simulation results shown in this figure can be found in Figure 5 - source data 1.

While Ca^2+^ uncaging stimuli are exquisitely suited to map the full range of synaptic responses, synaptic transmission is physiologically triggered by APs that induce short-lived Ca^2+^ transients. To stochastically predict responses to such time-varying Ca^2+^ stimuli we implemented our model using a Gillespie algorithm. After verifying that this model implementation agrees with the initial implementation (Figure 5 – figure supplement 1), we simulated responses to a typical AP-induced Ca^2+^ wave RRP SVs experience (Figure 5C, top panel). With 15 syts per SV, AP-induced EPSCs were large and synchronous, but reducing their number decreased response amplitudes (Figure 5C). Removal of one syt already reduced the average EPSC amplitude by ∼10% and removal of half (7/15) of its copies reduced it by ∼72% (Figure 5 – figure supplement 2, for representative example traces see Figure 5C). Note, however, that our model only describes the functioning of syt1/ syt2 and therefore does not include other Ca^2+^-sensors, like syt7 and Doc2B, which may mediate release in case of syt1/ syt2 loss (Bacaj *et al*., 2013; Kochubey *et al*., 2016; Kochubey and Schneggenburger, 2011; Luo and Sudhof, 2017; Sun *et al*., 2007; Wen et al., 2010; Yao et al., 2011).

As the number of syts per SV has a large impact on fusion kinetics we wondered to what extent fluctuations in the number of syts per SV affected the variance in AP-evoked responses in case of their imperfect sorting. Strikingly, however, varying the number of syts per SV over a large range (Poisson distribution with mean = 15, Figure 5D_1_,E, top panel) did not increase the variability of AP-evoked in synaptic responses while fluctuations of the RRP size largely impacted them (Figure 5D_2_,E). This shows that although release kinetics strongly depend on the average number of syts per SV, the system is rather insensitive to fluctuations around this number between individual SVs. Taken together, our data show that although only a subset of syts are required to simultaneously bind Ca^2+^ and PI(4,5)P_2_ to induce fusion, all SV syts contribute to the high rates of NT release by increasing the probability that several (i.e. three) syts simultaneously engage in dual Ca^2+^/PI(4,5)P_2_ binding.

Besides the number of syts, the PI(4,5)P_2_ levels also determine how likely it is for syts to engage in dual Ca^2+^/PI(4,5)P_2_ binding at an SV (see Figure 3 and Figure 2 – figure supplement 5). We therefore reasoned that upregulation of PI(4,5)P_2_ levels could potentially compensate for reduced syt expression. To investigate this, we refitted the models with reduced syt levels to the experimental Ca^2+^ uncaging data (Kochubey and Schneggenburger, 2011) and only allowed the PI(4,5)P_2_ concentration in the slots ([PI(4,5)P_2_]) to vary. Strikingly, increasing [PI(4,5)P_2_] fully rescued the characteristics of NT release upon reductions in syt levels down to 3 syts per SV (corresponding to an 80% reduction) by restoring number and speed of C2B domains engaging in Ca^2+^/PI(4,5)P_2_ dual binding (Figure 5 – figure supplement 2A-C). The required increase in [PI(4,5)P_2_] ranged from ∼1.1x (14 syts) to ∼10x (3 syts, Figure 5 – figure supplement 2D). These elevations also fully restored simulated AP-evoked responses when at least three syts per SV were present (Figure 5 – figure supplement 2E-F). Altogether, these data indicate that upregulating [PI(4,5)P_2_] is a potential, powerful compensatory mechanism to rescue reductions of NT release in case the number of (functional) syts per SV is reduced to no less than three. We note that this compensatory mechanism may complicate experimentally observed effects of stoichiometric changes.

### Evaluation of mutants affecting Ca^2+^ binding to the C2B domain reveals diverse effects on AP-evoked transmission

Ca^2+^ sensing of syts depends on negatively charged aspartate (D) sidechains of the C2B domain whose positions are optimal to bind Ca^2+^ ions (Fernandez *et al*., 2001). The local negative charges of the Ca^2+^ binding sites are reduced/neutralized upon Ca^2+^ binding. The Ca^2+^ binding pockets of the C2B domain have been extensively studied using various mutations (Bradberry et al., 2020; Guan et al., 2017; Kochubey and Schneggenburger, 2011; Lee *et al*., 2013; Mackler *et al*., 2002; Nishiki and Augustine, 2004; Shin et al., 2009). Mutations that remove or invert the negative charge of the Ca^2+^ binding sites (by mutation to asparagine (N) or lysine (K), ‘DN’ or ‘DK’) block Ca^2+^ binding and severely reduce exocytosis, even when co-expressed together with the wildtype protein (Bradberry *et al*., 2020; Kochubey and Schneggenburger, 2011; Lee *et al*., 2013; Mackler *et al*., 2002). Other mutations also interfere with Ca^2+^ binding and exocytosis but hold the same pocket charge (e.g. mutation to Glutamate, ‘DE’) (Bradberry *et al*., 2020). While both types of mutations may similarly interfere with Ca^2+^ binding, they may differentially affect the allosteric mechanism. The allosteric coupling between the Ca^2+^ and PI(4,5)P_2_ binding sites might be (in part) mediated by electrostatic interactions (van den Bogaart *et al*., 2012), which would imply that the negatively charged Ca^2+^ binding pocket repels PI(4,5)P_2_ until Ca^2+^ reverses the electrostatic charge, and vice versa. Following this assumption, charge altering mutations within the Ca^2+^ binding pockets (‘DN’, ‘DK’) would partially activate the allosteric coupling mechanism and thereby affect the domain’s PI(4,5)P_2_ affinity (which would not be the case in mutants conserving the charges (‘DE’)). We explored this in our model using two different hypothetical Ca^2+^ site mutants (i.e. mutations that do or do not affect PI(4,5)P_2_ affinity; Figure 6A). We investigated the effect of these mutants on AP-induced synaptic transmission under homozygous and heterozygous expression conditions (combined expression of mutant and WT with a total of 15 syts per SV; Figure 6B).

**Figure 6:**
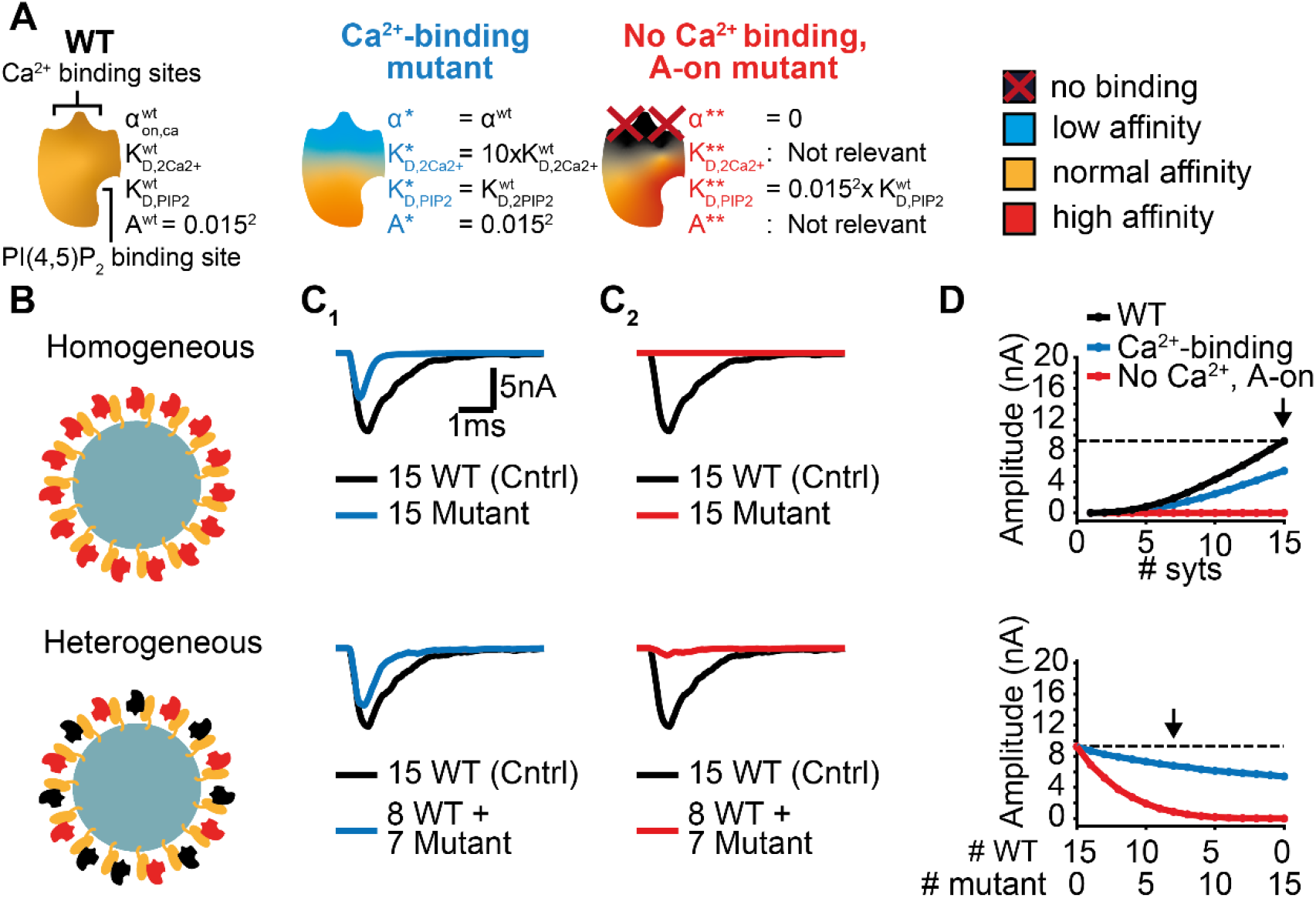
Systematic evaluation of the effect of mutant syts on simulated AP-evoked fusion. **A)** Illustration of a WT syt and two mutant syts. The “*Ca*^*2+*^*-binding”* mutant has a lower affinity for Ca^2+^ (K_D,2Ca2+_ 10x increased, i.e. β 10x increased). The “*no-Ca*^*2+*^ *binding, A-on”* mutant is not able to bind Ca^2+^ and has a high binding affinity for PI(4,5)P_2_, which is equal to the affinity for PI(4,5)P_2_ when the allostericity between Ca^2+^ and PI(4,5)P_2_ is “active” in WT syts (Ca^2+^-bound state). Because of the inability to bind Ca^2+^, allosteric interactions between Ca^2+^ and PI(4,5)P_2_ are not possible in this mutant. **B)** Illustration of homogeneous (top) and heterogeneous expression (bottom) of the mutants. Mutant syts are depicted in red, WT syts are depicted in black. **C)** Representative, stochastically simulated AP-evoked responses with homozygous (top, 15 mutant syt copies) and heterozygous (bottom, 8 WT and 7 mutant syt copies) expression of the different mutants (C_1_: “*Ca*^*2+*^*-binding”* mutant, in blue; C_2_: “*no Ca*^*2+*^ *binding, A-on”* mutant, in red). For each of the settings a representative trace of a condition with 15 WT syts is shown in black (control condition). A third mutation, the *“no Ca*^*2+*^ *binding, A-off”* was also explored (Figure 6 – figure supplement 1). **D)** Mean amplitudes of simulated AP-evoked responses (n=200) for the homogeneous (top), heterogeneous (bottom) of the different mutants, and WT syt (for homozygous condition only). Dotted line indicates the mean amplitude of simulated eEPSCs with 15 copies of WT syt (control). Arrow indicates the condition that is depicted in panels C. Simulation scripts can be found in Source code 1. Results from simulations can be found in Figure 6 - source data 1.

The first mutant, the “*Ca*^*2+*^*-binding”* mutant, had a lower Ca^2+^ affinity (10xK_D,2Ca2+_), but all other properties were the same as in the WT C2B domain. This mutant might be equivalent to mutation of the binding pocket which conserves its charge (e.g. ‘DE’). When homozygously expressed, this mutant showed eEPSCs with a ∼50% reduced amplitude and faster kinetics compared to the WT condition (Figure 6C_1_, Figure 6D top). Heterozygous expression (8 WT and 7 mutant syts per SV) only caused a small decrease in mean eEPSC amplitude compared to the expression of 15 WT syts per SV (Figure 6C_1_-D, bottom), showing that this mutant is relatively mild.

The second hypothetical mutation was designed to not only abolish Ca^2+^ binding, but to also mimic the Ca^2+^-bound state. Thereby this mutant featured a high PI(4,5)P_2_ affinity as if the allosteric interaction between Ca^2+^ and PI(4,5)P_2_ was permanently ‘on’. This might represent an extreme example of a mutation electrostatically reducing/inverting the negative charges of the Ca^2+^ binding pocket (e.g. ‘DN’, ‘DK’). We termed this mutant the “*no Ca*^*2+*^ *binding, A-on”* mutant (Figure 6A). This mutant showed no NT release in response to the Ca^2+^ transient in a homozygous condition (Figure 6C_2_-D, top), which is explained by its inability to bind Ca^2+^. A major detrimental effect of the mutant was observed when co-expressed with the wildtype protein: When half of the syts on the SV were mutated (heterozygote), the amplitude of simulated eEPSCs was strongly reduced (Figure 6C_2_-D, bottom). Merely four mutant proteins expressed together with 11 WT proteins already decreased eEPSC amplitudes by ∼70% (Figure 6D, bottom), indicating a strong dominant negative effect. The strong\ inhibition is a result of the mutant’s increased PI(4,5)P_2_ affinity leading to occupation of PI(4,5)P_2_ binding slots on the membrane with this Ca^2+^-insensitive mutant which blocks the association of the Ca^2+^ sensitive- and SV fusion promoting WT proteins. In comparison, a mutant not able to bind Ca^2+^ but having a normal PI(4,5)P_2_ affinity (“*no Ca*^*2+*^ *binding, A-off”* mutant, which could represent a more extreme form of the ‘DE’ mutant) had a much weaker effect (Figure 6 - figure supplement 1). This indicates that the allosteric interaction between Ca^2+^ and PI(4,5)P_2_ can play a prominent role in the severity of mutations.

### Rapid changes of accessible PI(4,5)P_2_ dramatically impact synaptic short-term plasticity

In our model we describe the PI(4,5)P_2_ levels in concentration units, because our model is based on syt affinities for Ca^2+^ and PI(4,5)P_2_ determined *in vitro* (van den Bogaart *et al*., 2012). The estimated concentration of PI(4,5)P_2_ not only depends on the local density of PI(4,5)P_2_ in the membrane, but also on the accessibility syt has to PI(4,5)P_2_. While all species (Ca^2+^, PI(4,5)P_2_, and syt C2B) are homogenously accessible in the aqueous solution of the *in vitro* setting (van den Bogaart *et al*., 2012), at the synapse the syt C2 domains have constrained motility due to their vesicular association and PI(4,5)P_2_ is restricted to (clusters on) the plasma membrane (Milosevic et al., 2005; van den Bogaart *et al*., 2011a). This implies that the positioning of SVs with respect to the plasma membrane has an impact on the PI(4,5)P_2_ concentration accessible to syts. We so far assumed that all syts of RRP SVs are exposed to the same PI(4,5)P_2_ levels (which could be the case if all vesicles are similarly docked to the plasma membrane). However, when considering more complex stimulation paradigms such as repetitive AP stimulation, this may no longer be valid as several studies reported activity-dependent changes in SV positioning on a millisecond timescale (Chang *et al*., 2018; Kusick et al., 2020; Miki et al., 2016). Such changes may thus contribute to short-term plasticity, the alteration of responses on a millisecond timescale (Abbott and Regehr, 2004; Kobbersmed et al., 2020; Neher and Brose, 2018; Silva et al., 2021). Recent studies reported that mutations of positively charged amino acids of the C2B domain (lysines, ‘Ks’, implicated in binding of PI(4,5)P_2_ and/or the SNAREs and arginines, ‘Rs’ implicated in binding the plasma membrane and/or the SNAREs resulted in a loss of SV docking and severely reduced neurotransmission (Chang *et al*., 2018; Chen *et al*., 2021; Li *et al*., 2006; Xue *et al*., 2008). Strikingly, SV docking in these mutants was rapidly restored within milliseconds after an AP which also led to enhanced synaptic transmission in response to a second AP given 10 ms after the first (Chang *et al*., 2018). We explored such a situation in the context of our model by driving exocytosis with two successive AP-induced Ca^2+^ transients and assuming either constant PI(4,5)P_2_ levels for syts in wildtype synapses (i.e. all RRP SVs similarly docked) or initially reduced and activity-dependent increasing PI(4,5)P_2_ levels for syts in mutant synapses (where SVs docked after the first AP) (Figure 7A). We studied the consequence of a mutation that would only affect SV docking at steady state (as may be the case upon mutation of the arginines 398 and 399 of mouse syt 1, ‘R398,399Q’) in (Figure 7). This resulted in a markedly decreased initial response (Figure 7B,C), but repeated activation induced a large facilitation of responses (indicated by a large paired pulse ratio: quotient of the second EPSC amplitudes divided by the first) (Figure 7D). Mutations of the C2B domain that reduce its PI(4,5)P_2_ affinity (as is likely the case upon mutation of the lysine residues 325 and 327 in syt1 or 327, 328 and 332 in syt2) may be more detrimental because even when the effective PI(4,5)P_2_ concentration accessible to syts is restored upon activity-dependent SV redocking, syt association to PI(4,5)P_2_ will still be less probable. We conclude that dynamic changes in the PI(4,5)P_2_ levels accessible to syts - which may be caused by activity dependent SV relocation – strongly impact synaptic short-term plasticity.

**Figure 7.**
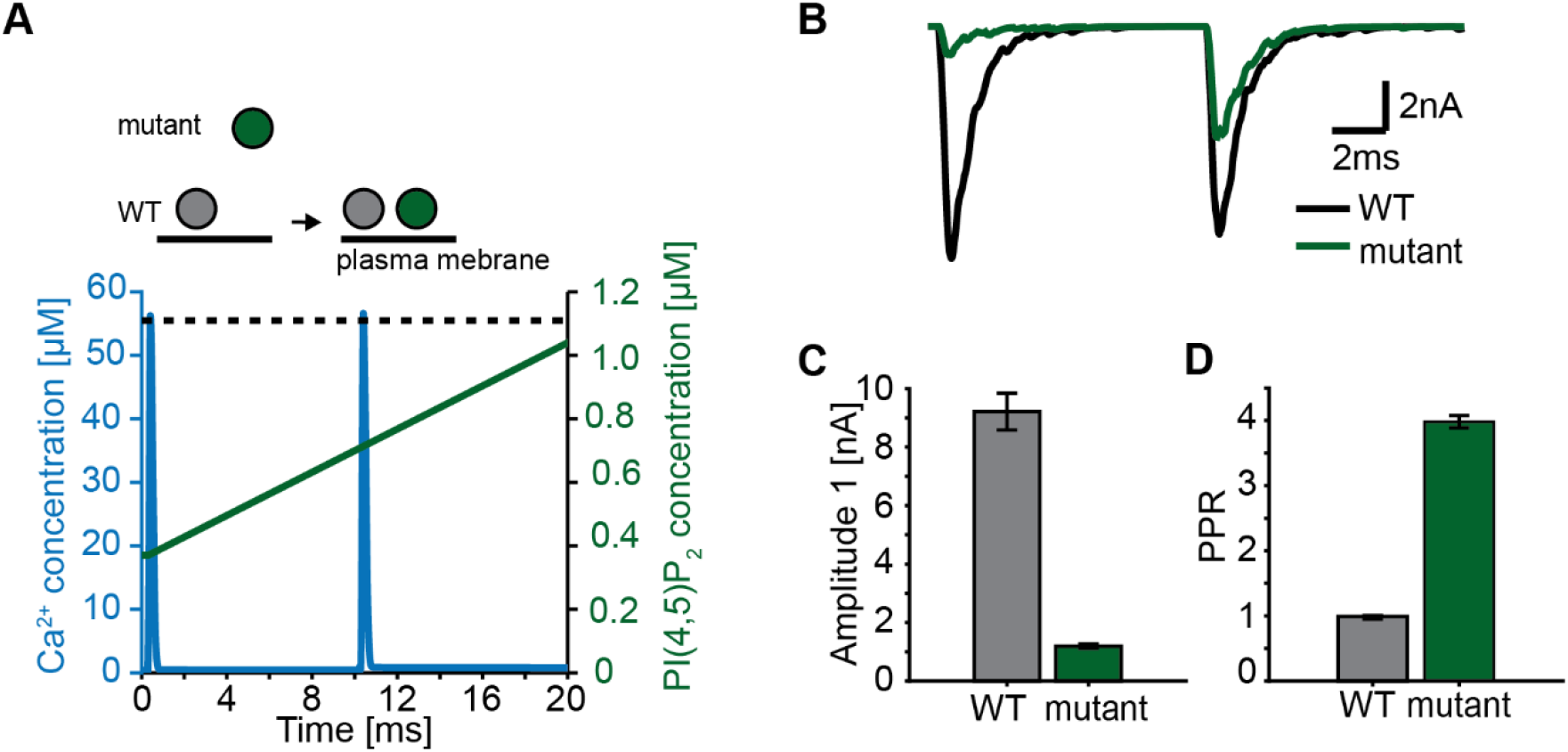
Paired-pulse stimulation in a membrane binding syt mutant. **A)** Time-course of [Ca^2+^]_i_ (blue) and [PI(4,5)P_2_](dashed black line: wildtype (WT), green line: mutant). Top panel illustrates the placement of vesicles with respect to the PM for SVs expressing WT syt (grey SVs) and SVs expressing a syt mutant deficient in membrane binding (green SVs, homozygous expression) before the first (left side of arrow) and second AP (right side of arrow). In WT conditions, most SVs reside close to the PM before the onset of the first stimulus. Before onset of the second pulse, WT SVs keep the same average distance to the PM. Mutant SVs, however, show a large distance to the PM at the onset of the first stimulus. Before the onset of the second AP, mutant vesicles move closer to the PM due to a Ca^2+^-dependent mechanism (Chang *et al*., 2018). The bottom panel shows the Ca^2+^ (blue) and PI(4,5)P_2_ (green) transients over time in a paired pulse stimulus (10 ms between stimuli). Due to the increased distance between the SV and the PM in the membrane-binding mutant, mutant vesicles are assumed to experience a lower [PI(4,5)P_2_] (solid green line) compared to WT SVs (dotted, black line) at the start of the first stimulus. Before the start of the second stimulus, mutant SVs move closer to the PM which increases the experienced [PI(4,5)P_2_] of these SVs. **B)** Representative eEPSCs simulated using the Ca^2+^- and PI(4,5)P_2_ transients depicted in A. **C)** Amplitude of the first eEPSC for WT and mutant. **D)** Paired-Pulse ratio (PPR) for WT and mutant. Data in C and D show mean ± SEM, using 50 repetitions and a variable RRP size (see Methods for details, the same RRP values were used for both the mutant and the WT condition). Simulation scripts can be found in Source code 1. Depicted simulation results can be found in Figure 7 - source data 1.

## Discussion

We here propose a quantitative, experiment-based model describing the function of syt in SV fusion on a molecular level based on biochemical properties determined *in vitro*. In our model, syt acts by lowering the energy barrier for SV fusion by dual binding to Ca^2+^ and PI(4,5)P_2_. When allowing at least three dual-bound syts per SV at a time, this model can explain the steep Ca^2+^ dependence of NT release observed at the calyx of Held synapse (Kochubey and Schneggenburger, 2011). Exploring this model led to the following conclusions:

1. The positive allosteric interaction between Ca^2+^ and PI(4,5)P_2_ is crucial for fast SV fusion as it stabilizes the dual-bound state which allows multiple syts to successively lower the energy barrier for SV fusion;
2. At least three syts must simultaneously engage in Ca^2+^/PI(4,5)P_2_ dual binding for fast SV fusion to achieve the speed and Ca^2+^ sensitivity inherent to synaptic transmission;
3. A high copy number of syts per SV ensures fast NT release by increasing the probability that three syts engage in Ca^2+^/PI(4,5)P_2_ dual binding;
4. Binding of syts to PI(4,5)P_2_ prior to the Ca^2+^ stimulus allows some SVs to fuse very fast (submillisecond).
5. The molecular resolution of this model can be used to study consequences of mutations.

### A syt-dependent switch on the energy barrier for SV fusion

Exocytosis is a highly energy-demanding reaction, for which the formation of the neuronal SNARE complex provides the energy (Jahn and Fasshauer, 2012). In our model we assume that syts regulate this process by lowering the activation energy barrier for exocytosis when they engage in Ca^2+^/PI(4,5)P_2_ dual binding. However, how Ca^2+^/PI(4,5)P_2_ dual binding to syt exactly reduces this energy barrier is not known. One possibility is that the energy is provided by the SNAREs themselves and that Ca^2+^/PI(4,5)P_2_ dual binding to syt relieves a clamping function, which syt itself or the auxiliary protein complexin exerts on the SNAREs (Courtney et al., 2019; Schupp *et al*., 2016; Tang *et al*., 2006; Zhou *et al*., 2015; Zhou *et al*., 2017). Alternatively – or additionally – syt’s dual binding Ca^2+^/PI(4,5)P_2_ might promote SNARE-mediated fusion by changing the local electrostatic environment (Ruiter *et al*., 2019; Shao et al., 1997). Furthermore, dual-binding syts could bring SVs closer to the plasma membrane, potentially below an upper limit for full SNARE complex assembly (Araç *et al*., 2006; Chang *et al*., 2018; Honigmann *et al*., 2013; Hui et al., 2011; Lin et al., 2014; Nyenhuis et al., 2019; van den Bogaart *et al*., 2011b; Xue *et al*., 2008). In line with these hypothetical working mechanisms, our estimated effect each dual-bound C2B domain has on the energy barrier height (4.85 k_b_T) is similar to the estimated energy barrier height for the final zippering step of the SNARE complex (Li et al., 2016). Syt’s Ca^2+^/PI(4,5)P_2_ dual binding might also promote fusion by inducing membrane curvature or favoring lipid rearrangement (Lai et al., 2011; Martens *et al*., 2007). In line with this reasoning, our estimated contribution of a syt engaging in Ca^2+^/PI(4,5)P_2_ dual binding is similar to estimates of syt1 membrane binding energies (Gruget *et al*., 2020; Gruget et al., 2018; Ma et al., 2017). In our model, we assume that multiple syts can simultaneously reduce the energy barrier for fusion. We here assumed that all syts exert the same effect on this energy barrier for fusion and that the effects of more dual-bound syts are additive. Whether or not this is the case will depend on the precise mechanism by which they shape the energy landscape. We show here that the simplest model (constant and independent contribution) is sufficient to reproduce the biological response.

Both the C2A and C2B domain of syt cooperate in SV exocytosis (Bowers and Reist, 2020; Gruget *et al*., 2020; Lee *et al*., 2013; Wu *et al*., 2021b). However, the exact role of the C2A domain in triggering SV fusion remains debated (Fernández-Chacón *et al*., 2002; Lee *et al*., 2013; Paddock et al., 2011; Sørensen et al., 2003; Stevens and Sullivan, 2003; Striegel et al., 2012). As mutation of the Ca^2+^ binding pockets of the C2A domain did not affect the affinities of Ca^2+^ and PI(4,5)P_2_ *in vitro* (van den Bogaart *et al*., 2012), we focused on the C2B domain in our model. This, however, does not imply that our C2B-domain-based model, which fully recapitulates the physiological responses of the synapse, does not indirectly describe properties of the C2A domain For instance, Ca^2+^ binding to the C2A domain may influence the Ca^2+^ affinity for the C2B domain (Sørensen *et al*., 2003),and this property may be collapsed into other parameters of model (e.g. our estimate of the PI(4,5)P_2_ concentration) that similarly determine this.

### Allostericity buys time to synchronize syts

Our modeling study proposes that the allostericity between Ca^2+^ and PI(4,5)P_2_ binding is essential for the syts to achieve fast, synchronous and sensitive NT release (van den Bogaart *et al*., 2012) (Figure 4 – figure supplement 1). With their experiment, Van den Bogaart and colleagues (2012) determined steady state affinities, which do not provide information on the association/dissociation rates. This means that the allosteric effect may either be due to speeding up the association or slowing down the dissociation of Ca^2+^/PI(4,5)P_2_ (Figure 1A). We here implemented the latter, a reduction of the unbinding rates of both Ca^2+^ and PI(4,5)P_2_ when both species were bound to the C2B domain, which leads to a stabilization of the dual-bound state. A stabilization of the Ca^2+^-bound state, which increases the lifetime of the state, was also essential to reproduce the Ca^2+^ dependence of release in the previously proposed five-site binding model (Heidelberger *et al*., 1994; Schneggenburger and Neher, 2000). We here show in the context of our model that increasing the lifetime of Ca^2+^/PI(4,5)P_2_ dual binding is particularly important to achieve fast fusion rates as it allows several C2B domains to simultaneously engage to lower the fusion barrier (Figure 4). The drawback of the strong allosteric interaction between the Ca^2+^ and PI(4,5)P_2_ bindings sites might be its potential involvement in the strong dominant-negative effects of some C2B domain mutations (Figure 6).

### The stoichiometry of the SV fusion machinery

Each SV contains multiple syt copies (Takamori *et al*., 2006), which can jointly participate in the fusion process. However, the number of syts that can simultaneously engage with PI(4,5)P_2_ located at the plasma membrane, and thus can cooperate during fusion, is likely limited. There are several possible explanations for this limit. First, the space between the vesicular and plasma membrane is limited and crowded by many synaptic proteins (Wilhelm et al., 2014). In addition, plasma membrane association of syt may require interaction with the SNAREs (de Wit *et al*., 2009; Mohrmann *et al*., 2013; Rickman and Davletov, 2003; Zhou *et al*., 2015), which limits the number of association points. Moreover, the inhomogeneous distribution of PI(4,5)P_2_ in the plasma membrane might put further constraints on association of syt to PI(4,5)P_2_ (Milosevic *et al*., 2005; van den Bogaart *et al*., 2011a). We found that at least three PI(4,5)P_2_ association sites (‘slots’) were required to explain the steep Ca^2+^ dependency of neurotransmitter release (Figure 2 A-C). These findings are compatible with a cryo-EM analysis that identified six protein complexes between docked SVs and plasma membrane (Radhakrishnan et al., 2021). Interestingly, irrespective of the number of slots, for models with three or more slots our analysis suggests that most fusion events at [Ca^2+^]_i_ > 1 *µ*M occurred after engaging three syts in Ca^2+^/PI(4,5)P_2_ dual binding (Figure 2 D-E). At lower [Ca^2+^]_i_ (0.5-1 *µ*M, Figure 2 - figure supplement 2B and Figure 2 – figure supplement 3B), the number of dual bindings leading to fusion was reduced to 1-2, indicating that higher [Ca^2+^]_i_ recruits additional syts to increase fusion rates.

The predicted number of three syts involved in fast exocytosis matches experimental estimates of the number of SNARE-complexes zippering for fast vesicle fusion (Arancillo et al., 2013; Mohrmann et al., 2010; Shi et al., 2012; Sinha et al., 2011) (but higher estimates in the number of SNARE complexes actively involved in fusion have also been reported (Wu et al., 2017)). Moreover, our model is consistent with a previous model of neurotransmitter release at the frog neuromuscular junctions that estimated that fusion is triggered by the binding of two Ca^2+^ ions to each of three syts. That model, which describes Ca^2+^ dynamics in the AZ in great detail, showed that many additional Ca^2+^ binding sites (20-40) were required to enhance fusion probability, because the probability of having a single Ca^2+^ molecule in the vicinity of SVs is extremely low (Dittrich et al., 2013). Similarly, in our model additional syts promote SV fusion by increasing the collision probability of syts with Ca^2+^ and PI(4,5)P_2_, and thereby increase the probability of (multiple) syts engaging in Ca^2+^/PI(4,5)P_2_ dual-binding for fast synaptic transmission (Figure 5). High protein abundance might also promote other collision-limited processes in fusion of SVs, and may provide an intuitive explanation for the high abundance of synaptobrevins on SVs (∼70 copies) which may need to assemble into SNARE complexes downstream of syt action (Takamori *et al*., 2006; van den Bogaart *et al*., 2011b).

Our model predicts that most RRP SVs have prebound PI(4,5)P_2_ via one syt at rest (Figure 3A), but it is the Ca^2+^-induced, allosteric PI(4,5)P_2_ affinity-increase that induces the dynamic assembly of three C2B domains in Ca^2+^/PI(4,5)P_2_ dual binding. This is fundamentally different from studies suggesting that 12-20 syts need to preassemble in higher-order complexes (rings) to execute their function in fusion (Rothman et al., 2017). A testable property to distinguish these possibilities is the sensitivity to reducing the number of syts per SV. If transmitter release relied on preassembled syt-rings, it would immediately break down if the number of syts was reduced to one preventing ring assembly, whereas our model predicts gradual effects (even for titration below *n*_*syt*_ =3) (Figure 5).

### Heterogeneity in PI(4,5)P_2_ concentration between different RRP SVs

The interaction between syt and PI(4,5)P_2_ has been shown to be essential in SV exocytosis (Bai *et al*., 2004; Li *et al*., 2006; Wu *et al*., 2021a), but also has been found to play a role in SV docking (Chang *et al*., 2018; Chen *et al*., 2021). Consistently, we observed that at resting synapses the majority of SVs (∼83%) contain at least one syt bound to PI(4,5)P_2_ in our model (Figure 3). The number of syts bound to PI(4,5)P_2_ per SV at rest highly influenced the release probability, leading to heterogeneity within the RRP (Figure 3) (Wolfel *et al*., 2007). As PI(4,5)P_2_ levels have a large impact on release kinetics (Figure 2 – figure supplement 5, Figure 5 – figure supplement 2), heterogeneity between RRP vesicles might further be enhanced by unequal PI(4,5)P_2_ levels.

In our model, we described PI(4,5)P_2_ levels in concentration units to constrain our model by using in vitro PI(4,5)P_2_ affinity measurements (van den Bogaart *et al*., 2012). However, this concentration does not only encompass the density of PI(4,5)P_2_ in the plasma membrane, but also includes the accessibility of syt to PI(4,5)P_2_. Several studies have shown that PI(4,5)P_2_ is distributed heterogeneously over the plasma membrane, in clusters that contain a high PI(4,5)P_2_ density (Honigmann *et al*., 2013; Milosevic *et al*., 2005; van den Bogaart *et al*., 2011a). Moreover, syts located closer to the plasma membrane will have increased access to PI(4,5)P_2_ (“see a higher PI(4,5)P_2_ concentration”) compared to those located further away. Taken together, this indicates that the PI(4,5)P_2_ concentration is likely to vary between RRP SVs and also between individual syts on the SV. Furthermore, this implies that once a syt has engaged in PI(4,5)P_2_ binding the successive engagement of additional syts might be favored for some (those facing towards the PM) and disfavored for others (those facing from the PM). While knowledge of these details could be helpful to construct a more realistic version of our molecular model, we currently do not possess the methodology to measure these properties. We therefore in our model simulated the simplest scenario where all syts have an equal probability of engaging in PI(4,5)P_2_ binding.

As the localization of syts with respect to the PM influences the accessibility of syt to PI(4,5)P_2_, mutations in synaptic proteins and stimulation protocols that alter SV docking will affect the PI(4,5)P_2_ concentration as it is implemented in our model (Chang *et al*., 2018; Chen *et al*., 2021; Kusick *et al*., 2020). Using a time-dependent PI(4,5)P_2_ concentration, we here illustrated the impact this might have on the short term plasticity of synaptic responses (Figure 7). This is a simplification, as we did not take the individual SV/syt distances to the plasma membrane into account. This distance is affected by several synaptic proteins, including syt1, Munc13 and Syt7 (Chen *et al*., 2021; Imig et al., 2014; Liu et al., 2016; Quade et al., 2019; Tawfik et al., 2021; Voleti et al., 2017). A role of these proteins in short term plasticity is firmly established, yet precise mechanistic details are still lacking (Jackman et al., 2016; Rosenmund et al., 2002; Shin et al., 2010). The extension of models based on molecular interactions such as presented here should allow for the recapitulation of more complex synaptic activity patterns relevant for neural processing. Particularly the molecular resolution of such models will be useful to conceptualize the importance of specific molecular interactions for physiological and pathological processes at the synapse.

## Materials and Methods

In this paper we propose a model for SV fusion induced by Ca^2+^ and PI(4,5)P_2_ binding to *n*_*syts*_ syts per SV, with at most *M*_*slots*_ syts per SV engaging in PI(4,5)P_2_ binding at the same time. We implemented the model in two ways for different simulation purposes: (1) an implementation based on the analytical solution of the model, and (2) an implementation following the Gillespie algorithm (Gillespie, 2007) (Matlab procedures for simulations can be found in Source code 1). In the first implementation we assume a constant [Ca^2+^]_i,_ (allowing us to simulate Ca^2+^ uncaging experiments: SV reactions at a Ca^2+^ level reached by the uncaging stimulus from a steady state starting point calculated for the resting Ca^2+^ concentration of 50 nM), whereas the second version was implemented to allow for [Ca^2+^]_i_ to vary over time (allowing us to simulate AP-evoked responses). Another important difference between the two approaches is that the analytical solution describes the binding state of an entire SV and the Gillespie version describes the binding state of each individual syt. Both implementations allow for stochastic evaluation of the model. The first implementation is used in Figure 2, 2 supplement 4, 2 supplement 5, 3, 3 supplement 1, 4C-D,F-G, 4 supplement 1, 5A-B, 5 supplement 1A-B. The second implementation is used to simulate the AP-evoked responses and individual SV binding states in Figures 2 supplement 2, 2 supplement 3, 4B,E, 5C-E, 5 supplement 1E-F, 6C-D, and 6 supplement 1, and 7B-D. Consistency between the two approaches was validated by comparison of simulation result distributions in quantile-quantile (Q-Q) plots (Figure 5 – figure supplement 2).

### SV states and possible reactions in the analytical version of the model

In the analytical solution of the model, we describe for each SV the number of syts having bound two Ca^2+^ ions, PI(4,5)P_2_, or both species. Since syts were assumed to work independently, their order is not relevant, and we therefore do not need to describe the binding state of each individual syt. The possible binding states of an SV are described in Figure 1 – figure supplement 2. Each state is represented by the triplet *(n,m,k)*, with *k* denoting the number syts having bound PI(4,5)P_2_, *m* denoting the number of syts having bound two Ca^2+^ ions, and *n* denoting the number of syts having bound both species and thereby having formed a dual binding. *n*_*syts*_ is the total number of syts per SV. *M*_*slots*_ restricts the number of syts having bound PI(4,5)P_2_ simultaneously (this includes syts having bound PI(4,5)P_2_ only (*k*), and those having formed a dual binding (*n*). Taken together, this implies that for all states in the model, it holds that

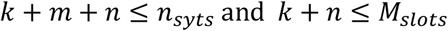

We numbered the states systematically following a lexicographic ordering, excluding the states that violate the inequalities described above. To illustrate, we write the ordering of all the states *(m,n,k)* with *n*_*syts*_*=3* and *M*_*slots*_*=2:*

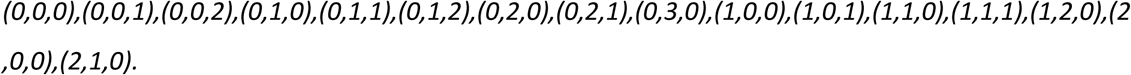

Besides these binding states, an additional state, *F*, describes whether the SV has fused. With *n*_*syts*_*=15* and *M*_*slots*_*=3*, a single SV in our model has 140 + 1 states. From any state, there are at most 9 possible reactions, one being SV fusion and the other 8 being (un)binding of Ca^2+^ or PI(4,5)P_2_ to/from a syt. The rates for the possible reactions of a single SV in this model are summarized in Table 3. In many cases, only a subset of the 8 (un)binding reactions are allowed because of the inequalities above (noted under “Condition” in the table).

**Table 3.**
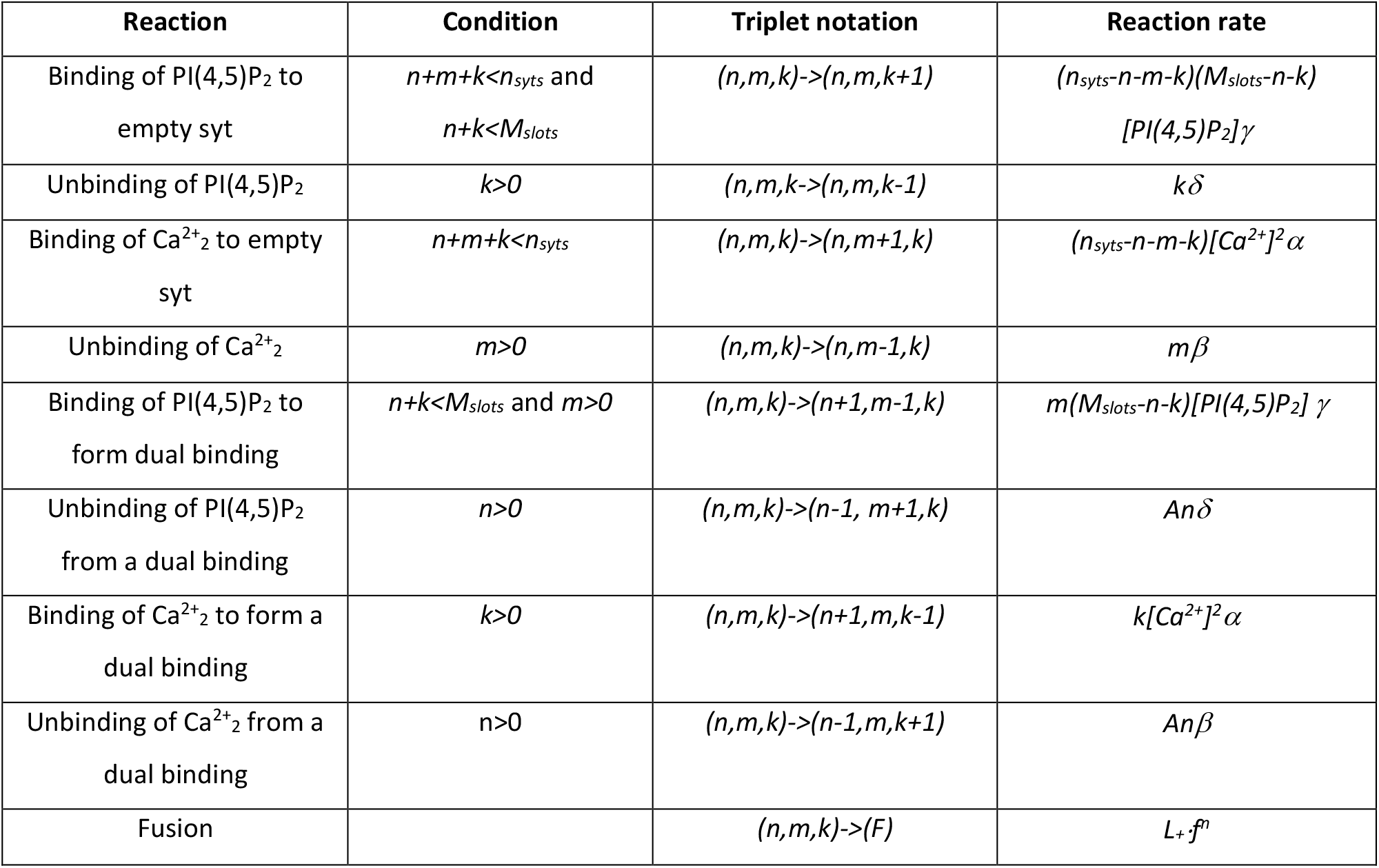
Overview of possible reactions and their rates in the model.

The reaction rates of (un)binding Ca^2+^ or PI(4,5)P_2_ are calculated as the number of syts available for (un)binding (computed using *n*_*syts*_, *n, m, k)* times the reaction rate constant (α, β, γ, δ), and, in the case of binding reactions, times the concentration of the ligand ([PI(4,5)P_2_] or [Ca^2+^]). We assumed binding of two Ca^2+^ ions to a single C2B domain. In our model this two-step process is simplified to a single reaction step by taking [Ca^2+^]_I_ to the power of two. This simplification is reasonable, because we assumed that syt could only associate to the vesicular membrane when two Ca^2+^ ions are bound, and binding of one Ca^2+^ ion would not induce an ‘intermediate’ association state to the membrane, nor would it affect the allosteric interaction. By simplifying the two Ca^2+^ binding/ubinding steps to one, we indirectly assumed high cooperativity between the two Ca^2+^ binding sites. To account for the limit on the number of syts bound to PI(4,5)P_2_, the number of available, empty slots, (*M*_*slots*_*-n-k)*, was multiplied on the PI(4,5)P_2_ binding rates. The fusion rate of the SV is computed by *L*_*+*_*·f*^*n*^ (similar to (Lou *et al*., 2005)), with *L*_*+*_ denoting the basal fusion rate and *f* the factor of increase in fusion rate by each dual binding being formed. *L*_*+*_ was set to 4.23e-4 s^-1^ to match the release rate measured at low [Ca^2+^]_i_, given an average size of the RRP of 4000 SVs (see below).

The affinities for Ca^2+^ and PI(4,5)P_2_ binding to syt were set to previously determined dissociation constants (*K*_*D,Ca2+*_*=β/α=221*^*2*^ *µM*^*2*^, *K*_*D,PI(4,5)P2*_*=δ/γ=20 µM)* obtained using *in vitro* microscale thermophoresis experiments (van den Bogaart *et al*., 2012). For determination of the dissociation constant of Ca^2+^, van den Bogaart and colleagues assumed binding of a single Ca^2+^ ion to the C2AB domain (van den Bogaart et al., 2012). The K_D,2Ca2+_ of the reaction describing binding of two Ca^2+^ ions, in our model, can be computed from the experimentally derived dissociation constant by taking it to the power of two. This was corroborated by re-fitting the experimental data with a hill coefficient of 2, which yielded a similar K_D,2Ca2+_ value of ∼221^2^ (data not shown).

The *in vitro* experiments revealed a change in syt1 Ca^2+^ affinity upon binding PI(4,5)P_2_, and vice versa (van den Bogaart et al., 2012), indicating a positive allosteric relationship between the two species. We assumed this allosteric effect was due to a stabilization of the dual-bound state by lowering of the unbinding rates of Ca^2+^ and PI(4,5)P_2_ with a factor (A=(3.3/221)^2^ =0.00022) and occurs when both species have bound. Upon dual binding, both rate constants for unbinding Ca^2 +^ and PI(4,5)P_2_ are multiplied by A, since any closed chemical system must obey microscopic reversibility (Colquhoun et al., 2004). Using the biochemically defined affinities, the number of free parameters in our model was constrained to:

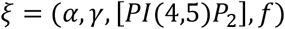

The values of *β* and *δ* were determined according to the affinities for each choice of *α* and *γ*.

### The steady state of the system

The steady state of the system before stimulation was determined at a resting, global [Ca^2+^]_i_ of 0.05 µM (except for simulations with Ca^2+^ levels below this basal value, for those we assumed [Ca^2+^]_rest_=[Ca^2+^]_i_). To compute the steady state, we assumed that no fusion took place, ignoring the very low fusion rate at resting [Ca^2+^]_i_. Under these conditions the model is a closed system of recurrent states and obeys microscopic reversibility, i.e. for every closed loop state diagram, the product of the rate constants around the loop is the same in both directions (Colquhoun *et al*., 2004). Microscopic reversibility implies detailed balance, meaning that every reaction is in equilibrium at steady state. Thus, for any two states S_i_ and S_j_ which are connected by a reaction, the steady state distribution obeys

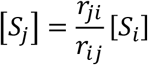

where [S_i_] and [S_j_] are steady state quantities and r_ij_ and r_ji_ are the reaction rates between S_i_ and S_j_. Using this property, we calculated the steady state iteratively by setting the population of the first state (state (0,0,0)), to 1, and thereafter iteratively computing the population of the following state (following the lexicographic ordering as described above) using the following formula’s:

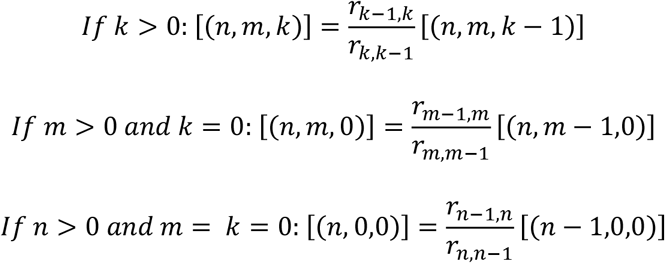

Afterwards, each state was divided by the sum of all state values and multiplied by the number of SVs in the RRP. In our model simulations, the size of the RRP was variable and followed a gamma distribution with a mean of 4000 SVs and a standard deviation of 2000 SVs, based on experimental estimates from the calyx of Held (Wölfel and Schneggenburger, 2003). In the following calculations we use *φ* to denote the steady state probability vector (i.e. normalised the sum to 1).

### Computation of fusion probabilities and fusion rate

The analytical implementation of our model allowed us to compute the fusion rate and cumulative fusion probabilities with a constant [Ca^2+^]_i_ after stimulus onset (t=0), thereby mimicking conditions in Ca^2+^ uncaging experiments. The constant [Ca^2+^]_i_ makes the model a homogenous Markov Model. The transition rates of the model can be organized in the intensity matrix, *Q*, such that,

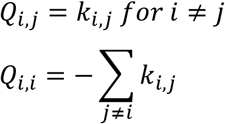

where *k*_*i,j*_ is the rate of the reaction from state *i* to state *j*. Given initial conditions, *φ* (steady state normalized to a probability vector), free model parameters, *ξ*, and calcium concentration, *C*,

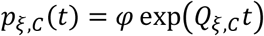

is the distribution of SV states a time t, i.e., a *1* x *n*_*states*_ vector with element *i* being the probability of being in state *i* at time point *t*. The single SV cumulative fusion probability is the last element, which we will denote with a subscript F,

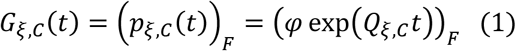

The fusion rate of a single SV can be calculated directly as the last element of the derivative of (1):

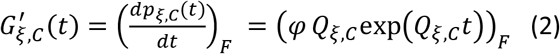

Multiplying (1) and (2) with the number of SVs yields the cumulative fusion function and fusion rate function, respectively. For simulation of peak release rates and release latencies (in Figure 2, 4, 5), we computed (1) and (2) using the best fit parameters from fitting with M_slots_=3 and n_syts_=15. This was done for 31 [Ca^2+^]_i_ values ranging from 0.001 µM to 80 µM (0.001, 0.1, 0.2, …., 0.9, 1, 1.25, 1.5, 1.75, 2, 2.5, 3, 4, …, 9, 10, 20, 30, …, 80 µM) for *t*∈[0,100] ms with a time step of 0.01 ms. In addition, the functions were calculated in the same way using the best fit parameters for M_slots_=1,2,4,5,6 with n_syts_=15 for simulations depicted in Figure 2A-C. In some conditions, especially at low [Ca^2+^]_i_, a longer span of the cumulative fusion probability was required and was calculated with the same time step size.

### Computation of peak release rates

The peak of the fusion rate can be computed by multiplying the maximum value of the single SV fusion rate function, (2), with *n*_*ves*_. To allow for a variable RRP size, a set of 1000 *n*_*ves*_ values were drawn according to the RRP size distribution, the peak release rates were determined, and the mean and 95 % prediction interval determined (Figure 2A, 2 supplement 4, 2 supplement 5, 4 supplement 1, 5A, 5 supplement 1A) for each Ca^2+^ concentration.

For parameter exploration (Figure 2G) and for computing the release rates in the fitting routine, it was not feasible to calculate the fusion rate over 100 ms with high temporal precision. Instead, we implemented a custom search algorithm (scripts can be found in accompanying zip-file “Source_code1.zip”), which was constructed to shorten calculation time by taking advantage of the release rate function being unimodal. We first found a time point, *t*_*max*_, at which 75-90 % of the SVs had fused. Having computed different time points in the time interval *[0,t*_*max*_], gave us an interval in which the fusion rate showed a local maximum. The algorithm then narrowed the time interval down until a time of peak was found with a precision of 0.01 ms. This method shortened simulation time considerably.

### Stochastic simulation of release latencies

Release latencies (Figure 2A, 2 supplement 4, 2 supplement 5, 4 supplement 1, 5A, 5 supplement 1A), which are defined as the time point of the fifth SV fusion event after the onset of simulation, were simulated stochastically by drawing *p*_*i*_∈(0,1), *i = 1,…,n*_*ves*_, from the uniform distribution for each of the 1000 repetitions and each evaluated [Ca^2+^]_i_. Each of these random numbers corresponds to the fusion time of an SV, which can be determined using the single SV cumulative fusion probability function (1). To obtain the time point of the fifth SV fusion, the fifth lowest *p*_*i*_ was selected. The corresponding time of fusion was found by interpolation of the single SV cumulative fusion probability, (1). In the corresponding figures, the medians were plotted, since the probability distribution of the release latencies (derived below) was skewed, and the reported data points were single measurements.

### Fitting the model to experimental data

We next fitted the model to already published data describing the Ca^2+^-dependence of NT release in the mouse calyx of Held (Kochubey and Schneggenburger, 2011). The data consist of measurements from Ca^2+^ uncaging experiments, where the release latency, defined as the time point of the fifth SV fusion event, and the peak release rate were estimated at different [Ca^2+^]_i_. Besides the four free model parameters, ξ, an additional parameter, *d*, was fitted. *d* is a constant added to the release latencies computed by the model to account for the experimentally observed delay (Kochubey and Schneggenburger, 2011).

Since the variance in the experimental data points also contains information on the underlying biological mechanism, we wanted to take the distribution of individual data points into account when obtaining estimates of the unknown parameters. We therefore derived the likelihood function, which describes how well the model captures the distribution of the release latencies. Obtaining this function for the peak release rates was not feasible. The experimental peak release rates were therefore compared to the average model prediction. Both measures of describing the correspondence between model simulations and experimental data were combined in a cost value which was optimized to estimate the best fit parameters (the lower this cost value the better the correspondence between model predictions and experimental data).

The best fit was obtained by minimizing the following cost function:

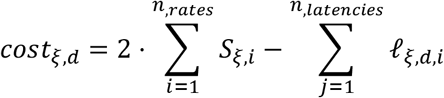

where *i=1,…, n*_*rates*_ and *j=1,…, n*_*latencies*_ are the indices of the experimentally measured [Ca^2+^]_i_,

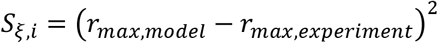

are the squared deviations of the peak release rates (1/ms^2^) and *ℓ* is the logarithm of the likelihood of the release latencies (see derivation below). To combine the two measures of distance between model and experimental data, the squared deviation of the peak release rates was multiplied by a factor 2 before subtracting the logarithm of the likelihood of the release latencies. The cost value was minimized using the inbuilt Matlab function *fminsearch*, which uses a Nelder-Mead simplex method. *fminsearch* was run with different initial parameter values to verify that the global minimum of the cost function was found. During the fitting, the lower and upper bounds of *d* were set to, respectively, 0.3 ms and 0.405 ms (with the upper bound corresponding to the smallest release latency in the experimental data set). α,γ, and [PI(4,5)P_2_] had an upper bound of 10^10^, and all free parameters needed to be positive. The maximum number of iterations and function evaluations was set to 5000.

### The likelihood function of release latencies with fixed RRP size

To fit the model to the experimental release latency measurements, we derived the likelihood function, which is the probability density function of the model for given parameters evaluated at the experimental data points. Thus, optimizing the likelihood function yields parameters for which the data points are most likely if the model is true. We first derive the likelihood of release latency in the case of a fixed RRP size (*n*_*ves*_).

We define the stochastic variable *X*, which describes the stochastic process of the state of a single SV in our model. The fusion time of the SV, *τ*, is defined as

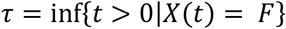

where *τ* itself is a stochastic variable. We define a stochastic vector, *Z*, which consists of all *n*_*ves*_ fusion time points in a single experiment. They come from independent, identically distributed stochastic processes with cumulative distribution function *G*_*ξ,C*_*(t)*, given in (1). As the release latency is defined as the time of the fifth SV fusion, we order the *Z* variable outcomes (*z*_*(1)*_, *z*_*(2)*_, *…, z*_*(nves)*_) from first to last fusion time. Using the transformation

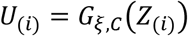

we obtain a sequence of stochastic variables, *U*_*(i)*_, which are uniformly distributed on the interval (0,1). The ordering is preserved, since *G*_*ξ,C*_*(t)* is monotonically increasing, and *U* has probability density function

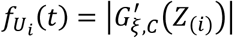

with respect to *t. G*_*ξ,C*_*’(t)≥0* is given in (2). From order statistics it follows that the *k*^*th*^ ordered *U* is beta-distributed with probability density

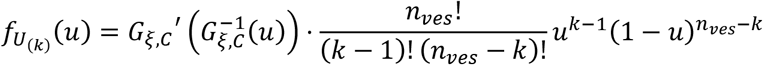

where *u = G*_*ξ,C*_*(t)*. Thus, the transformed variable *U*_*(k)*_, is beta-distributed, with

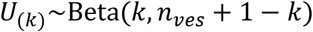

In the case of the release latency, we are interested in the fifth fusion event (*k=5*). Thus, with a fixed RRP size, the likelihood value for the release latency observations, *T*^*^=(t^*^_1_, t^*^_2_, …, t^*^_M_), at all *M* Ca^2+^ concentrations is

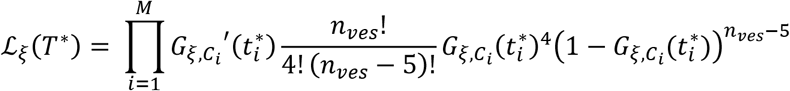

In the optimization we minimize minus the log-likelihood:

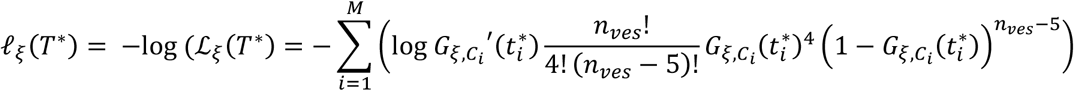

which is equivalent to maximizing the likelihood function.

### The likelihood of release latencies with variable RRP size

In our model, the RRP size is assumed to follow a Gamma distribution. Let *x* denote the RRP size, *f*_*RRP*_*(x)* the probability density of the Gamma distribution, and *u= G*_*ξ,C*_*(t)*. The probability density of the release latency at Ca^2+^ concentration *C*_*i*_ is given by

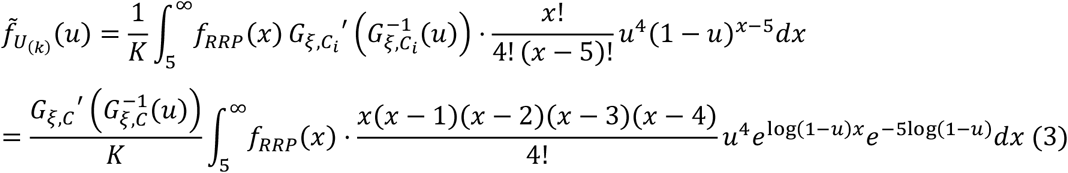

where *K* is a normalization constant, K=1-P(x<5)≈1. The lower limit of the integral reflects that the release latency is only defined when there are more than 5 SVs in the RRP. In simulations this corresponds to redrawing the RRP size whenever an RRP size < 5 SVs occurs, which happens with probability ∼3e-11, and is accounted for in the normalization constant K in the following. Inserting the probability density function of a Gamma distribution with shape parameter *k* and scale parameter *θ*, we get:

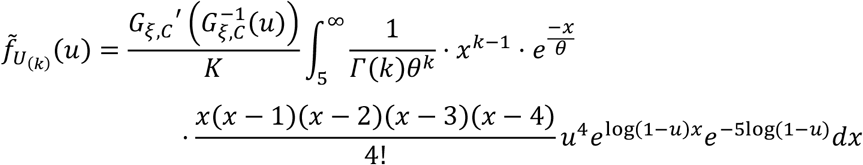

We now define the following variables:

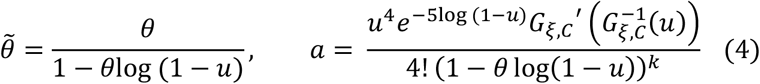

By factoring out and substituting in the above equation we get

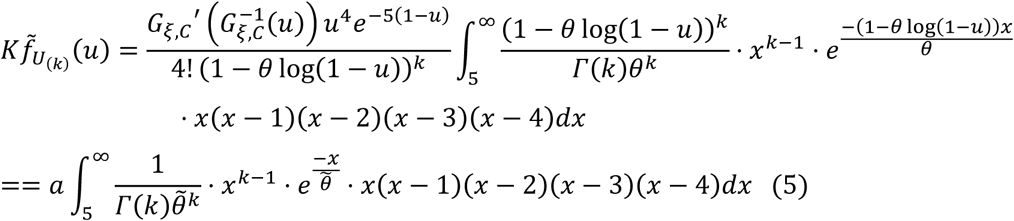

Furthermore, we have

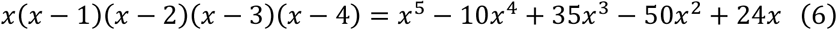

Since

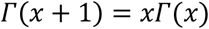

we can derive the following useful formula:

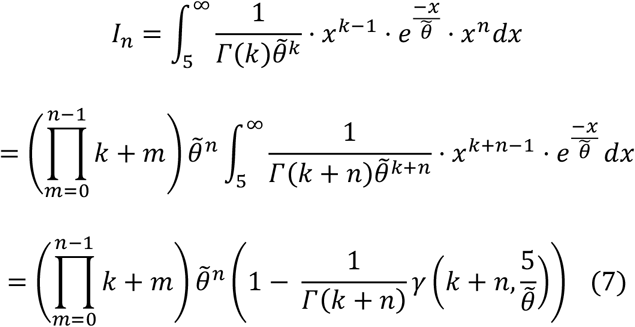

The third equality follows from the fact that the function in the second integral from above is the probability density function of a gamma distribution with shape parameter *k+n* and scale parameter 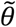. We therefore replace it with the cumulative distribution function of the gamma distribution, where *γ* is the lower incomplete gamma function. Combining (4), (5), (6), and (7) yields an explicit expression of the likelihood of a single delay in (3), as

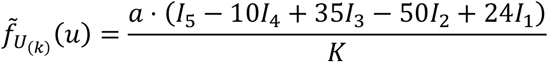

with

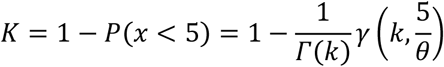

We then minimize the sum of minus the log-likelihoods of the release latency observations.

### Syt binding states in the Gillespie simulation of model

In the Gillespie algorithm, the binding state of each individual syt is tracked. The state of the system at time point *t, X(t)*, is given by a *n*_*syt*_ x *n*_*ves*_ matrix. Each element in this matrix describes the binding state of an individual syt using a 2-digit coding system; 00 for no species bound to syt, 01 for PI(4,5)P_2_ bound, 10 for two Ca^2+^ ions bound, and 11 for both species bound (dual-binding syt). As with the analytical implementation, each syt can undergo in total 8 different (un)binding reactions (Figure 1A), depending on the binding state of the respective syt. The fusion rate, which depends on the number of dual-bound syts per SV, is determined for the entire SV.

### Determining the initial state of the system

The steady state (initial state, *X(0)*) was computed using the same method as described above (see section ‘*The steady state of the system’*) using [Ca^2+^]_i_ = 0.05 µM as the resting condition. This resulted in *φ*, the probability vector of a single SV to be in the different SV-states at steady state. To stochastically determine *X(0)*, we first determined the binding state for each SV, *i*.*e*. how many dual bindings are formed (*n)* and how many syts have bound Ca^2+^ (*m)* and how many PI(4,5)P_2_ (*k)*. For that we drew *p*_*j*_∈(0,1), *j = 1…,n*_*ves*_, from the uniform distribution. The state number of the *j*^*th*^ SV, *s*, was determined by:

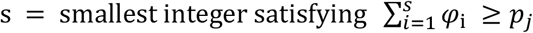

Via the ordering of states explained above, *s* can be linked to the state triplet *(n*_*s*_,*m*_*s*_,*k*_*s*_). As the order of syts is irrelevant for model simulation, this information on the state of SV_j_ was transferred to the *j*^th^ column of the *X(0)* matrix in a systematic way: The first *n*_*s*_ elements were labeled with ‘11’; elements *n*_*s*_*+1* to *n*_*s*_*+m*_*s*_ were labeled with ‘10’; and elements *n*_*s*_*+m*_*s*_*+1* to *n*_*s*_*+m*_*s*_*+k*_*s*_ were labeled with ‘01’, and the remaining elements *(n*_*s*_*+m*_*s*_*+k*_*s*_*+1):n*_*syt*_ were set to ‘00’.

### Gillespie algorithm-based simulations of the model

For stochastic evaluation of the model by the Gillespie algorithm (Gillespie, 2007), we next introduced the propensity function *b*, which is defined by:

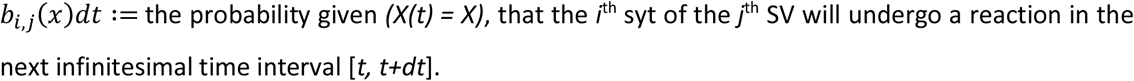

For element *i,j* in *b*, the total reaction propensity is the sum of propensities of possible reactions given the binding state of the syt and can be computed as follow:

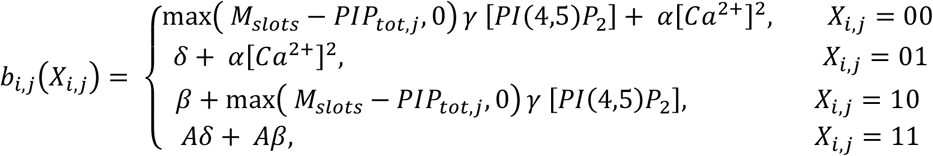

with *PIP*_*tot,j*_ the total number of syts of SV *j* bound to PI(4,5)P_2_. To include the propensity for fusion of an SV, an additional row in *b, b*_*f*_, describes the propensity of fusion for each SV in the matrix;

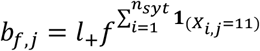

This makes *b* a matrix of *(n*_*syt*_*+1)* × *n*_*ves*_. We denote the sum of all elements in *b* by *b*_*0*_. Using *b*_*0*_ and 3 random numbers (r_n_ ∈(0,1), *n = 1,2,3)* drawn from the uniform distribution, we determined the time step to the next reaction and which SV and syt this reaction affects. The time to next reaction, *τ*, is given by

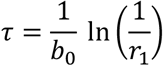

since it is exponentially distributed with rate *b*_*0*_. The index, *j*, of the SV undergoing a reaction is the first index that satisfies:

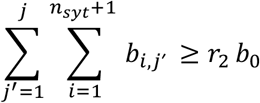

Similarly, the index *i* of the syt in SV *j* undergoing a reaction is the smallest integer fulfilling:

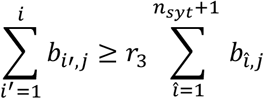

If the row index *i* equals *n*_*syt*_+1, a fusion reaction occurs. The fusion time (*t + τ*) is saved and all the propensities of SV *j* in *b* are set to 0. If *i* is smaller or equal to *n*_*syt*_ a binding or unbinding reaction of one of the two species occurs. To determine which of the four possible reactions is occurring, we define an additional propensity matrix, *b*_*react*_. The first element in *b*_*react*_ denotes the propensity of PI(4,5)P_2_ binding, the second element the propensity of Ca^2+^ binding, and the third and fourth element the unbinding of PI(4,5)P_2_ and Ca^2+^, respectively. These elements are given by:

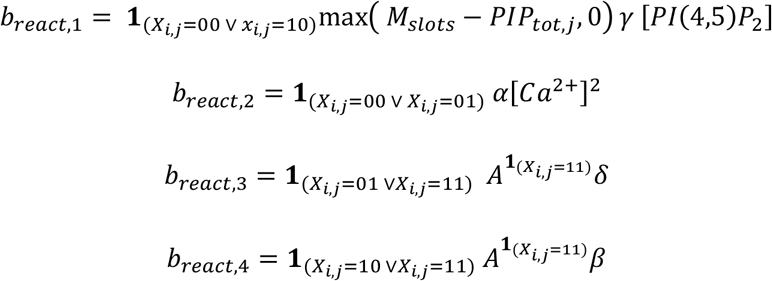

Additionally, we define the transition matrix *V*, which describes the change in the state of *X*_*i,j*_ induced by the four reactions:

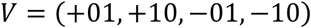

A fourth random number, r_4_ ∈(0,1), drawn from the uniform distribution, determines which reaction occurs via:

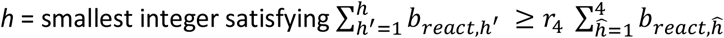

The state of the corresponding SV and syt, *X*_*i,j*,_ is replaced by *X*_*i, j*_ + *V*_*h*_ and t by *t* + *τ*. Then *b* is updated according to the change in *X*, and all steps are repeated. This iterative process continues until all SVs are fused, when simulating the model with a fixed [Ca^2+^]_i_.

When simulating AP-evoked responses (Figure 5, Figure 6), we used a Ca^2+^ transient describing the microdomain [Ca^2+^]_i_ sensed locally by primed SVs in the mouse calyx of Held upon AP stimulation (Wang *et al*., 2008). This Ca^2+^ transient also formed the basis for the Ca^2+^ signal used to simulate a paired pulse stimulus (Figure 7), where the transients were placed with a 10 ms interval. Additionally, for the paired pulse stimulus, we added a residual Ca^2+^-transient to the signal (exponential decay with amplitude: 0.4 µM, decay time constant: 0.154 s^-1^). Similar to the uncaging simulations, the [Ca^2+^]_i_ before the onset of the stimulus was 0.05µM. Since the Ca^2+^ concentration is a factor in the reaction rates, the propensity matrices *b* and *b*_*react*_ were not only updated to the new state matrix, *X(t* + *τ*), but also to a new [Ca^2+^]_i_ at time *t* + *τ*. [Ca^2+^] at time *t* + *τ* was determined from the Ca^2+^ transient using linear interpolation, and *b* and *b*_*react*_ were updated correspondingly. As the propensity of Ca^2+^ binding is largely dependent on [Ca^2+^]_i_, the time step between two updates in [Ca^2+^]_curr_ and the propensity matrices was set to be at most 8e^-4^s. If *τ* determined from *b* at time *t* was larger than this time step, no reaction occurred, but the system and [Ca^2+^]_curr_ were updated to time *t*+8e^-4^s. The model was evaluated until the end of the Ca^2+^ transient. Similarly, in the case of a variable PI(4,5)P_2_ transient (Figure 7), the PI(4,5)P_2_ concentration was updated at least every 8e^-4^s. Because this approach requires simulation of all individual (un)binding reactions and fusion events it is not feasible to perform 1000 repetitions. Instead, simulations were repeated 200 times (Figure 5, 6). Like with the computation of the release latencies and maximal fusion rates, we assumed a variable RRP. However, instead of drawing random pool sizes from a gamma distribution, we used the 200 quantiles of the pdf of the RRP sizes, because of the limited number of repetitions and the large impact of the RRP size on the model predictions. In Figure 7, we reduced the number of repetitions to 50, to represent an experimental condition more closely. The RRP was drawn randomly once. This set of random RRP values was used for both the mutant and the WT condition displayed in the figure.

### Simulating the model with mutant syts

For mutations in syt that affect the binding and unbinding rates of PI(4,5)P_2_ and Ca^2+^, the procedure described above was repeated with adjusted parameters when simulating a model containing only mutant syts. For a model in which mutant proteins were expressed together with WT syts (simulations of heterozygous condition in Figure 6), the procedure was changed slightly.

For a model with *p* WT and *q* mutant syts, the number of states of an SV increases drastically and was now described by six values; the number of WT syts bound to Ca^2+^, PI(4,5)P_2_ or both, and the number of mutant syts bound to Ca^2+^, PI(4,5)P_2_ or both. The dimensions of the Q-matrix used to compute the steady-state probability of a single SV increased proportionally with *n*_*states*_. *X(t=0)* was computed using the same principle as described above, with the important difference that the first *p* rows represented the binding status of the WT syts, and row *p+1:p+q* that of the mutants. In *b* this ordering of WT and mutant syts is the same. The parameters used to compute b_react_ depended on whether a reaction occurred to a WT syt, *i* ≤ *p*, or a mutant syt, *n*_*syt*_ ≥ *i* > *p*.

### Simulation of EPSCs

Simulated EPSCs were obtained using both model implementations. The analytical implementation of our model was used to simulate fusion times for a constant [Ca^2+^]_i_ (Figure 4D,G). The Gillespie version of the model was used to simulate AP-evoked EPSCs with or without mutant syts (Figure 5C-E, Figure 5 - figure supplement 1E-F, Figure 6C-D, 6 supplement 1, and 7B-D-E). In both approaches the stochastically determined fusion times, determined as described above, were rounded up to the next 0.02 ms, leading to a sampling rate of 50 kHz. The sampled fusion times were convolved with a mEPSC to generate simulated EPSCs. The standard mEPSC used for deconvolution followed the equation described by Neher and Sakaba (Neher and Sakaba, 2001):

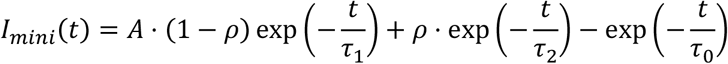

with *τ*_1_ = 0.12 ms (time constant of fast decay), *τ*_2_ = 13 ms (time constant of slow decay), *τ*_0_ = 0.12 ms (time constant of rise phase), *ρ* = 1.e-5 (proportion of slow phase in decay), and *A* being a normalization constant making the amplitude 60 pA. Parameter values were chosen such that the kinetics of the mEPSC would match events measured in the Calyx of Held (Chang et al., 2015). In Figure 4D,G traces show three randomly chosen eEPSCs in each panel. Representative eEPSC traces shown in Figures 5, 6, 6 supplement 1, and 7 are simulated eEPSCs with the amplitude closest to the mean eEPSC amplitude.

### Simulating AP-evoked EPSCs with variable syt

To investigate the effect of variability in the number of syts expressed per SV on variance between simulated AP-evoked traces model (Figure 5), we first had to determine the steady state. For this we computed the probability vector of a single SV to be in the different SV-states at steady state (*φ*) for n_syt_=1,…,50. Subsequently, for each SV and each repetition (n=1000) a random number of n_syt_ was drawn from the Poisson distribution with mean=15. When the value 0 was drawn, it was replaced by n_syt_=1. Using these values and *φ* determined for each n_syt_, we computed the steady state matrix (X(0)) as described above (“Determining the initial state of the system”). To reduce computation time, we evaluated a model containing 100 vesicles 40 times instead of evaluating 4000 vesicles simultaneously. The fusion times obtained when driving the model with the Ca^2+^-transient were combined afterwards. This is valid since all SVs act independently in the model. For the condition with a variable RRP size, fusion times were selected randomly until the RRP size of that specific repetition was reached. Afterwards, the fusion times were convolved with the mEPSC to obtain simulated AP-evoked responses. To quantify the variance in the traces, we computed the standard deviation of the simulated eEPSCs at each data point (300 data points corresponding to the time interval 0-6ms) and summed those values.

## Acknowledgements

We would like to thank Ralf Schneggenburger and Holger Taschenberger for providing the experimental Ca^2+^ uncaging dataset and AP-induced Ca^2+^ transient. This work was supported by grants from the Deutsche Forschungsgemeinschaft (Emmy Noether Programme, Project Number 261020751 and the TRR 186, Project Number 278001972 to A.M.W), the Novo Nordisk Foundation (NNF20OC0062958 to S.D.; NNF19OC0058298 to J.B.S and a Young Investigator Award NNF19OC0056047 to A.M.W.), the Lundbeck Foundation (R277-2018-802 to J.B.S.) and the Independent Research Fund Denmark (8020-00228A to J.B.S.).

## Additional files

Supplementary files:

- “Source_code1.zip”: MATLAB scripts used for simulations.
- Transparent reporting form
- Source data files for all figures and figure supplements

## Supplementary figures

**Figure 1 – figure supplement 1:**
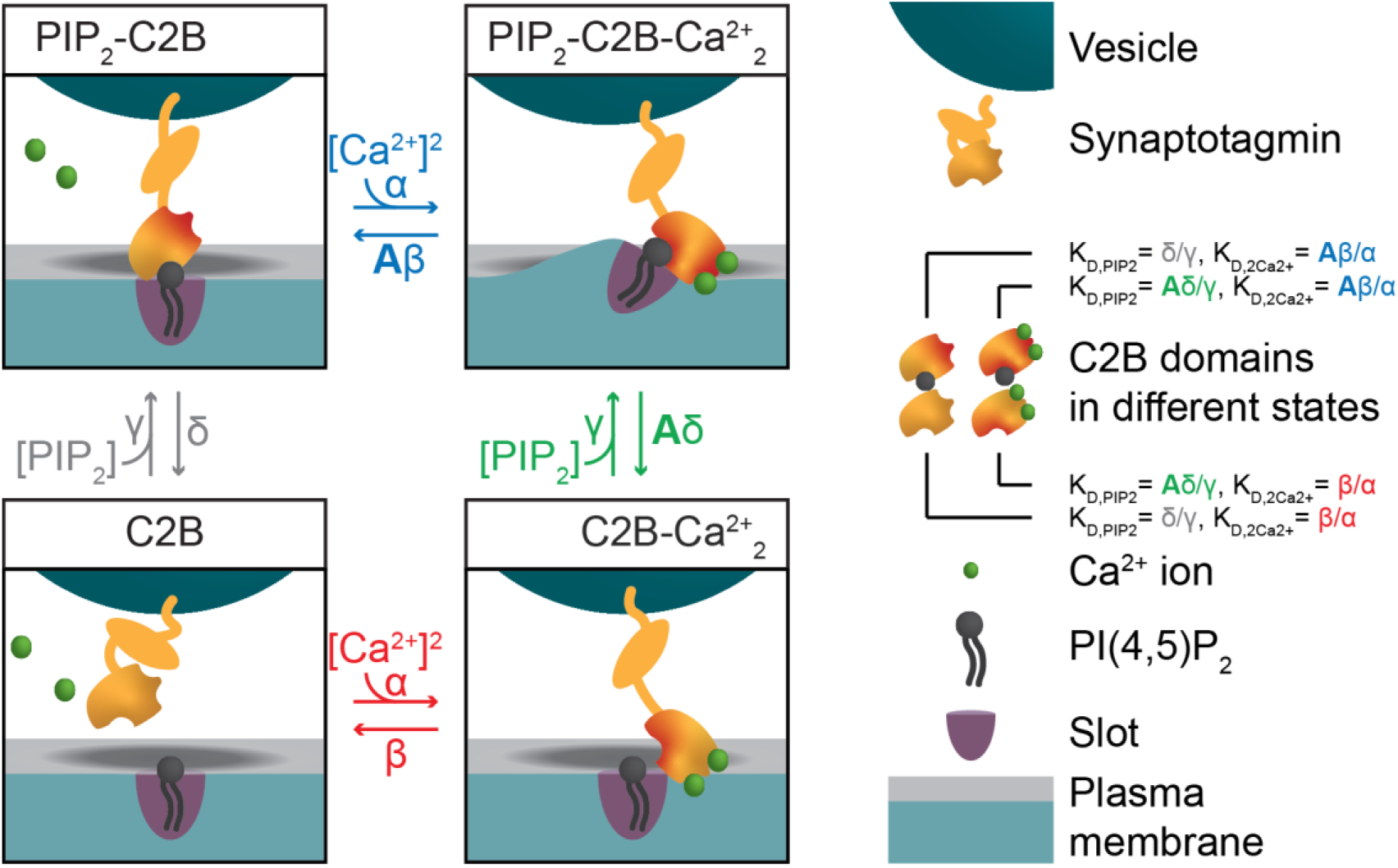
Alternative reaction scheme of a single syt in which Ca^2+^ binding leads to association to the plasma membrane. Reaction scheme of a single syt C2B domain alternative to the one shown in Figure 1A. Instead of an association to the vesicular membrane, Ca^2+^ binding to the C2B domain in this illustration leads to an association with the plasma membrane. In this illustration, the dual-bound state affects the curvature of plasma membrane. Our model does not make a distinction between either of the two illustrations.

**Figure 1 – figure supplement 2:**
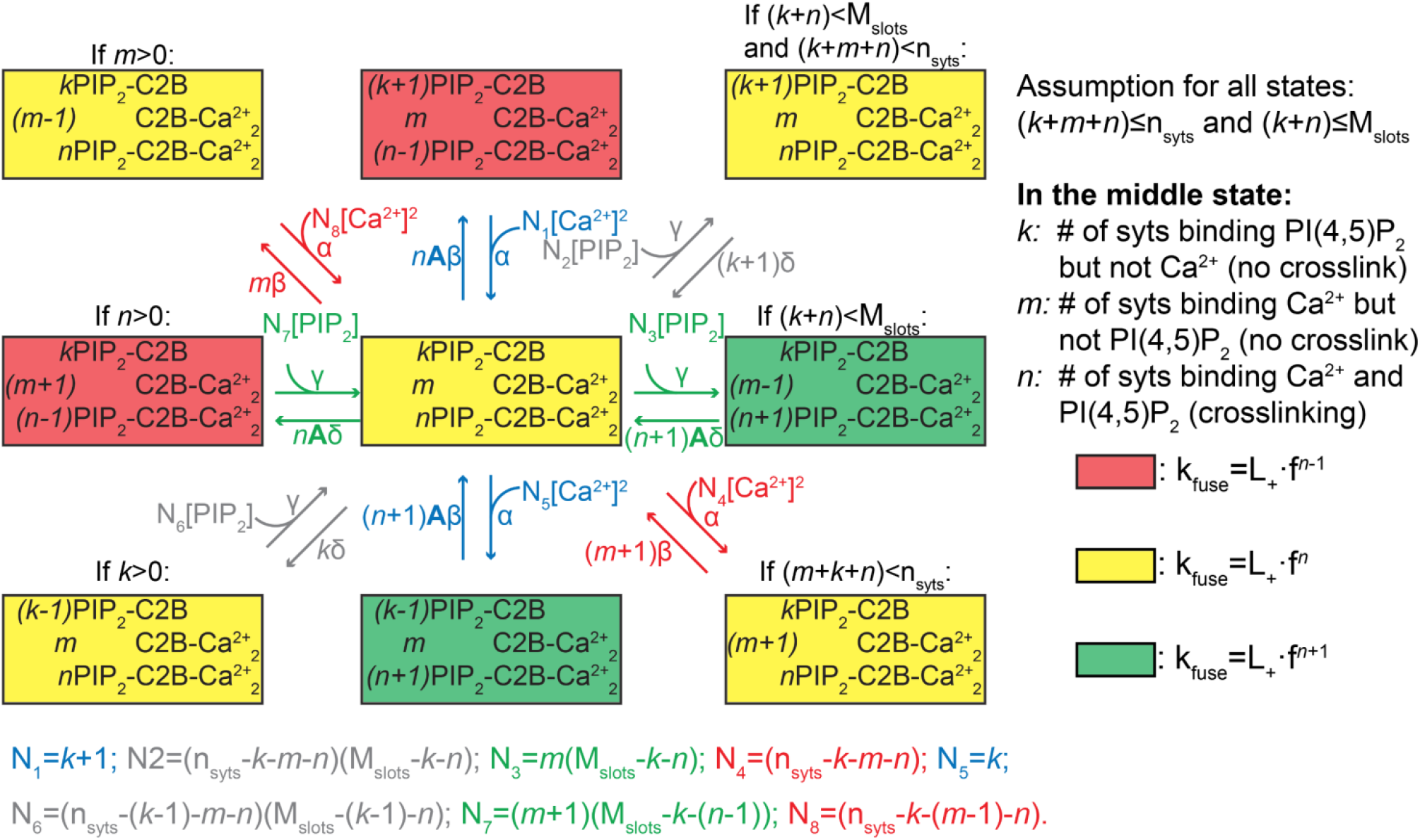
Reaction scheme for all reactions of an entire SV. The diagram shows the possible (un)binding reactions and indicates the relative fusion rate of an SV with *n*_*syts*_ syts and *M*_*slots*_ slots. The state in the center of the diagram represents an SV with *k* syts having bound only PI(4,5)P_2_, *m* syts having bound only two Ca^2+^, and *n* syts having formed dual bindings by binding both PI(4,5)P_2_ and two Ca^2+^. From this state, the SV can (un)bind Ca^2+^ or PI(4,5)P_2_ (unbinding moves the SV to the state to the left or up, binding to the right or down). The number of syts, *n*_*syts*_, and the number of slots, *M*_*slots*_, limit the possible number of syts engaging in ligand binding in general and in binding of PI(4,5)P_2_, respectively, which is indicated by the assumption in the top right corner. Whether the SV can undergo a specific reaction (that is, whether a certain state exists and the SV can transit to it) depends on its current binding state as indicated by the assumptions above some of the states. The color shading of the states indicates the value of the fusion rate relative to the other states in the diagram. Yellow shading means a fusion rate equal to that from the state in the center. Red and green shading means a factor *f* lower and higher fusion rate, respectively. For detailed description of the reaction rate equations, see <u>Methods</u>.

**Figure 2 – figure supplement 1:**
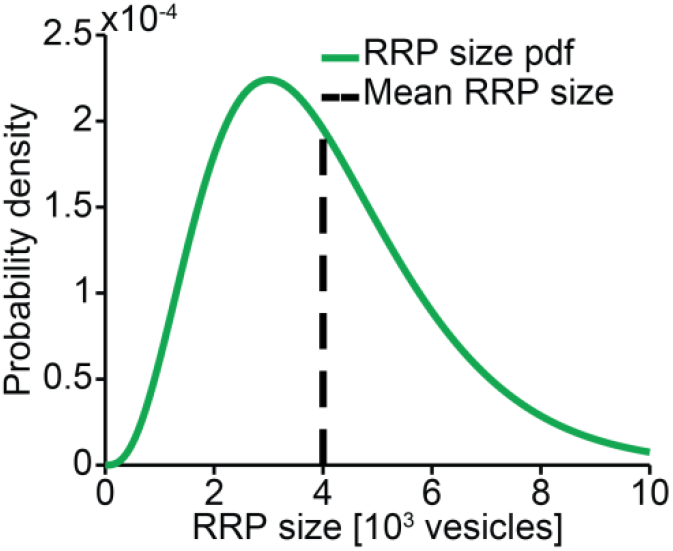
RRP distribution. We assumed the RRP size to follow a gamma distribution with mean 4000 and SD 2000.

**Figure 2 – figure supplement 2:**
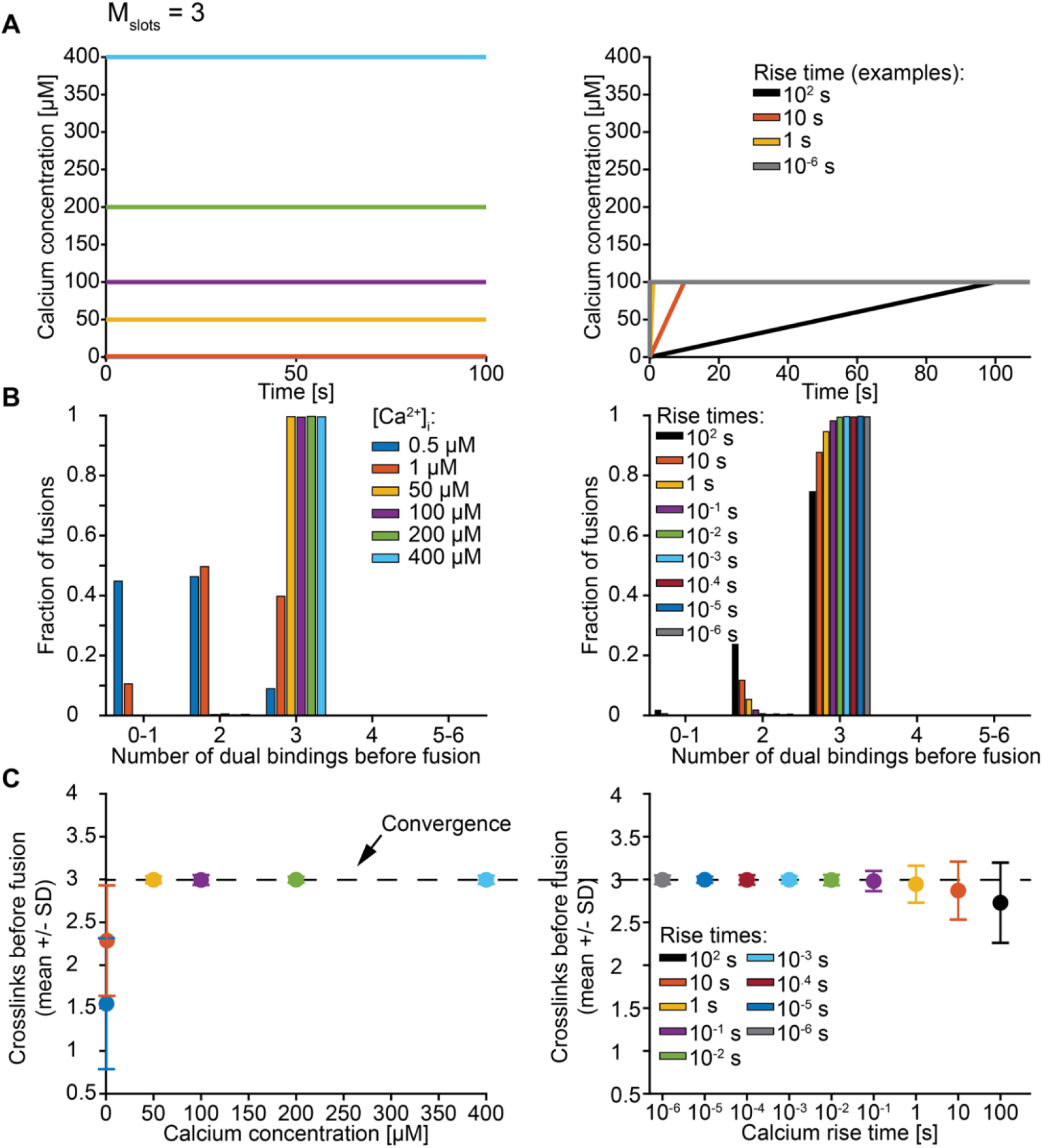
Exploration of the number of dual bindings formed before fusion of an SV with M_slots_=3. **A)** The Ca^2+^ signal used in simulations with step (left) and ramp (right) Ca^2+^ functions. In simulations with step Ca^2+^ the concentrations rises instantly at t=0 from the basal Ca^2+^ of 50 nM to various constant concentrations. In simulations with ramp Ca^2+^ the Ca^2+^ concentration increases linearly from the basal concentration of 50 nM to 100 µM Ca^2+^ with various rise times. **B)** The number of dual bindings formed before fusion for various Ca^2+^ concentrations (step Ca^2+^, left) or Ca^2+^ rise times (ramp Ca^2+^, right) as depicted in A. The bars show proportion of 10000 stochastically simulated SVs. The number of dual bindings formed before fusion increased with increasing step Ca^2+^ concentrations and decreasing Ca^2+^ rise times. At high concentrations or fast rise times, most fusions took place after forming 3 dual bindings. **C)** Average number of dual bindings formed before fusion in simulations of 10000 SVs with Ca^2+^ signals as depicted in A. The average number of dual bindings formed before fusion increases with increasing step Ca^2+^ concentration and decreases with increasing Ca^2+^ rise time. A plateau is reached at an average of 3 dual bindings at high step Ca^2+^ concentrations and low rise times. Error bars show ± standard deviation.

**Figure 2 – figure supplement 3:**
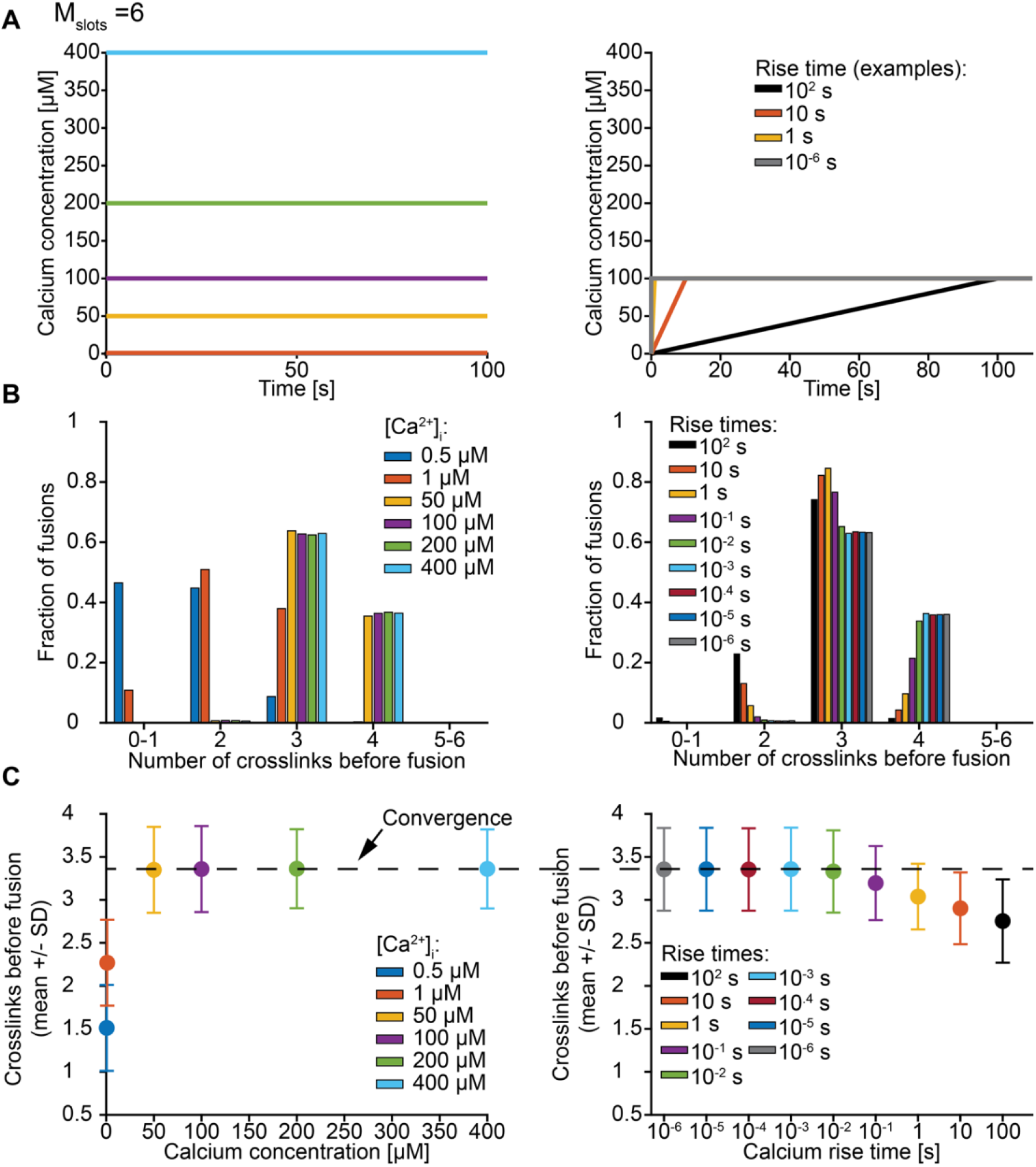
Exploration of the number of dual bindings formed before fusion of an SV with M_slots_=6. **A)** The Ca^2+^ signal used in simulations with step (left) and ramp (right) Ca^2+^ functions. In simulations with step Ca^2+^ the concentrations rises instantly at t=0 from the basal Ca^2+^ of 50 nM to various constant concentrations. In simulations with ramp Ca^2+^ the Ca^2+^ concentration increases linearly from the basal concentration of 50 nM to 100 µM Ca^2+^ with various rise times. **B)** The number of dual bindings formed before fusion for various Ca^2+^ concentrations (step Ca^2+^, left) or Ca^2+^ rise times (ramp Ca^2+^, right) as depicted in A. The bars show percentages of 10000 stochastically simulated SVs. The number of dual bindings formed before fusion increased with increasing step Ca^2+^ concentrations and decreasing Ca^2+^ rise times. At high concentrations or fast rise times, most fusions took place after forming 3-4 dual bindings. **C)** Average number of dual bindings formed before fusion in simulations of 10000 SVs with Ca^2+^ signals as depicted in A. The average number of dual bindings formed before fusion increases with increasing step Ca^2+^ concentration and decreases with increasing Ca^2+^ rise time. A plateau is reached at an average of 3.4 dual bindings at high step Ca^2+^ concentrations and low rise times. Error bars show ± standard deviation.

**Figure 2 – figure supplement 4:**
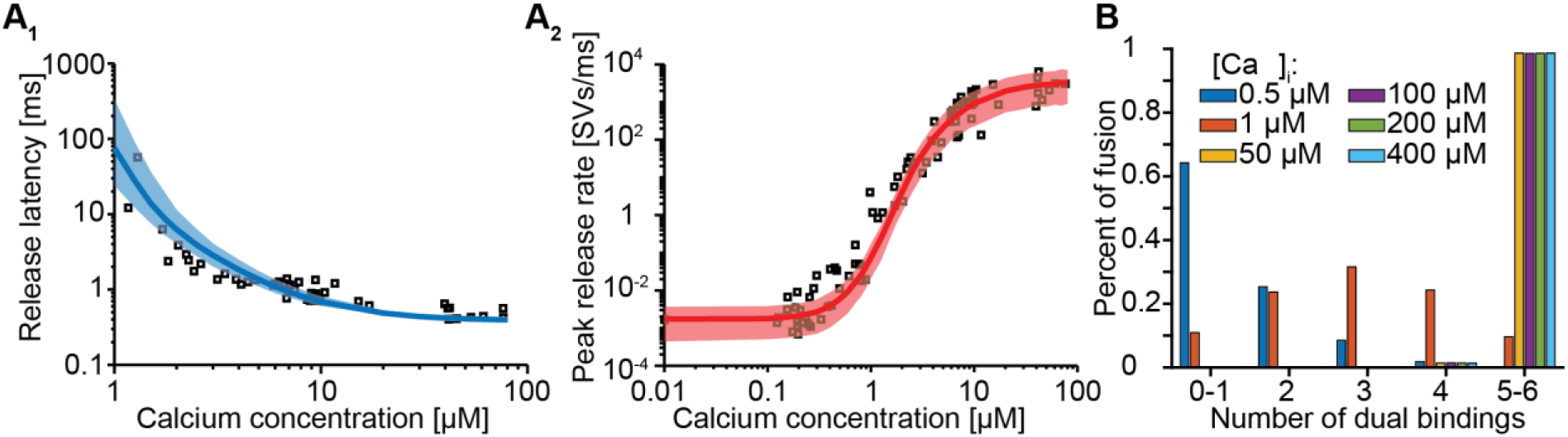
Forcing fusion from a state in which 5-6 syts are dual-binding Ca^2+^ and PI(4,5)P_2_ causes a too steep Ca^2+^ dependecy of the peak release rates. **A)** Best fit results for a model with 6 slots forced to show most fusion events when 5/6 dual-bounds were formed. We achieved this by setting *f* to a fixed value of 12.79 (computed following: 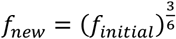, with *f*_*initial*_ being the best fit value at M_slots_=6 of *f*_*initial*_: 163.5) and we only fitted the four remaining parameters. The best fit results show larger deviation from the experimantally obtained release latencies (A_1_) and peak release rates (A_2_) compared to the initial fit with (*M*_*slots*_=6 and *f* as a free parameter). Solid lines show median release latencies (A_1_) and mean peak release rates (A_2_) predicted by the model from 1000 repetitions per simulated [Ca^2+^]_i_. The shaded areas indicate the 95% prediction interval of the model. **B)** Number of dual bindings formed before fusion for various Ca^2+^ concentrations. The bars show the proportion of 10000 stochastically simulated SVs that fuse with a certain number of dual bounds formed. Most fusion events occur when 5-6 dual bounds have formed. Best fit parameters: α=22.96 µM^-2^s^-1^, γ=0.49 µM^-1^s^-1^, [PI(4,5)P_2_]=12.71 µM, f=12.79 (fixed), d=0.3850 ms. Experimental data points in the figures are replotted from Kochubey and Schneggenburger (2011).

**Figure 2 – figure supplement 5:**
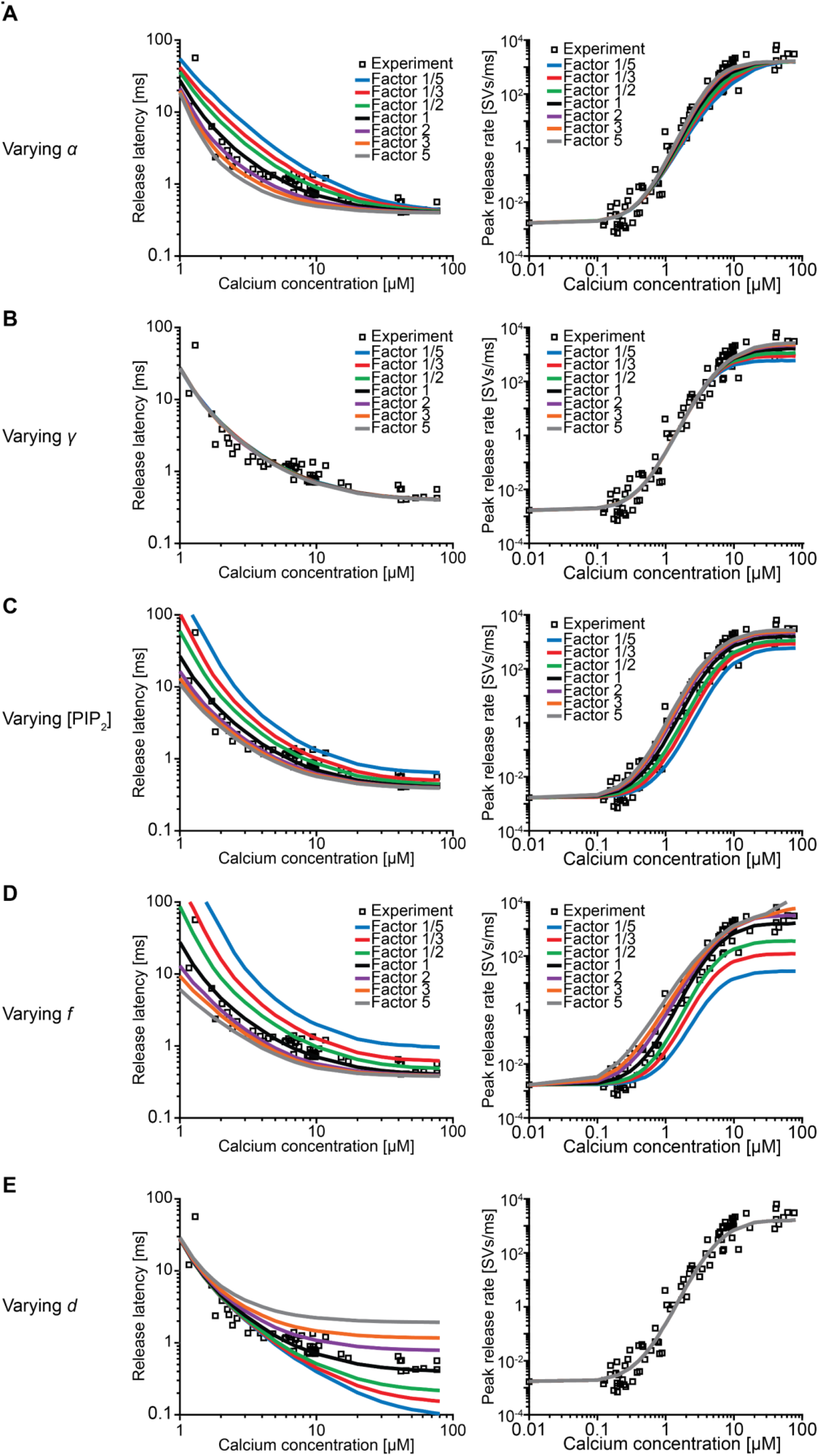
Effect of the free parameters on release latencies and peak release rates. **A-E)** Effect on release latencies (left) and peak release rates (right) when varying *α* (A), *γ* (B), [PI(4,5)P_2_] (C), *f* (D), and *d* (E). The best-fit values of each of the parameters were increased and decreased by a factor 2, 3 and 5. Varying α and γ leads to a change in β and δ so that the measured affinities for both Ca^2+^ and PI(4,5)P_2_ are still matched (van den Bogaart *et al*., 2012). (**A)** When varying *α* the release latency is mostly affected in the middle range of [Ca^2+^]_i_. The effect on the peak release rates is very small **B)** Varying *γ* has no effect on release latencies. The effect of varying γ on peak release is larger at high [Ca^2+^]_i_ compared to low [Ca^2+^]_i_ reflecting that at high [Ca^2+^]_i_ PI(4,5)P_2_ binding limits dual binding formation. **C)** Varying [PI(4,5)P_2_] has a large effect on the release latencies and peak release rates. The effect of varying [PI(4,5)P_2_] is much larger compared to varying γ, as changing [PI(4,5)P_2_] only affects the binding rate of PI(4,5)P_2_, whereas changing γ leads to an equal change in δ to match the published affinity values. Furthermore [PI(4,5)P_2_] affects the steady state distribution. **D)** Varying *f* has a large effect on both release latency and fusion rate, as it directly impacts the effect of dual binding formation on SV fusion rates. **E)** Varying the added delay, *d*, only shifts the release latencies linearly and does not affect the peak release rates. Experimental data points in these figures are replotted from Kochubey and Schneggenburger (Kochubey and Schneggenburger, 2011).

**Figure 3 – figure supplement 1.**
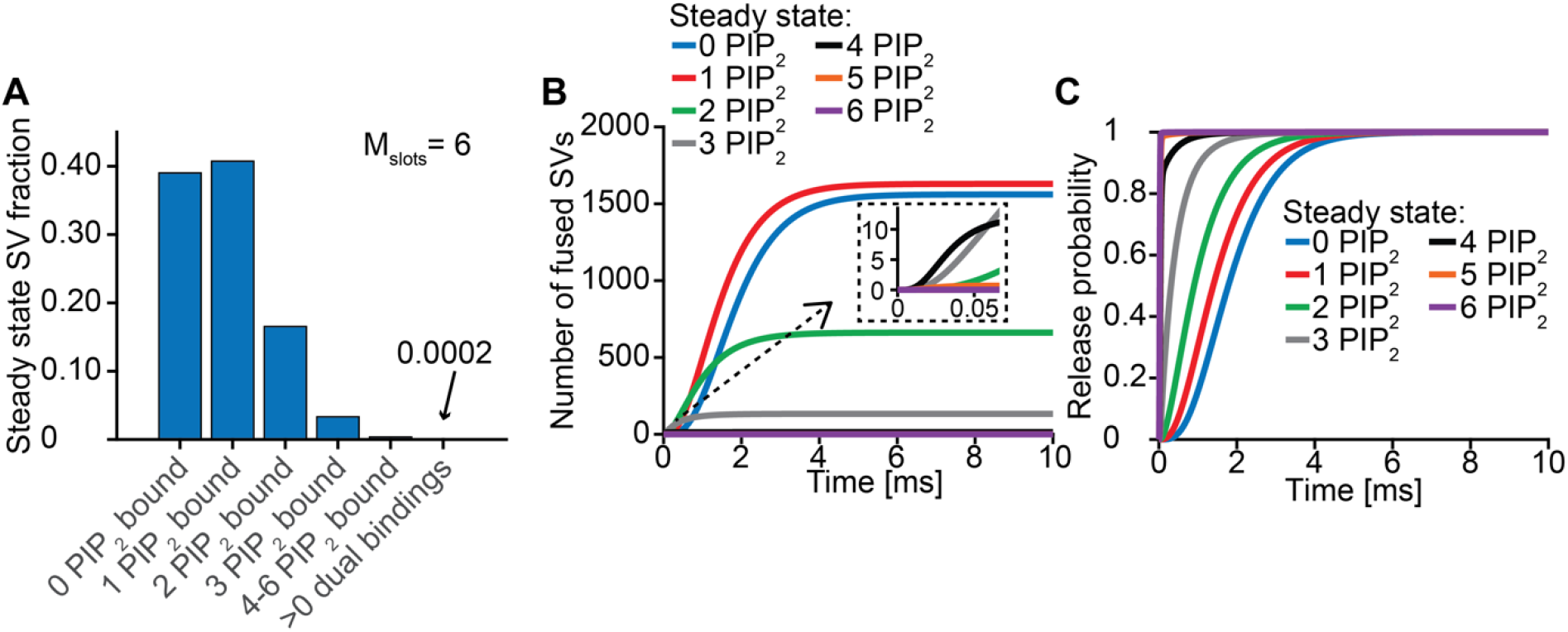
Syt-binding to PI(4,5)P_2_ prior to Ca^2+^ stimulus underlies very fast SV fusion (model with M_slots_=6). **A)** PI(4,5)P_2_ binding status of SVs at steady state. At resting [Ca^2+^]_i_ of 50 nM, more than 40 % of SVs have bound a single PI(4,5)P_2_ molecule (not including those that have formed a dual binding), more than 15 % has bound two PI(4,5)P_2_, while less than 5 % has bound three PI(4,5)P_2_. Very few SVs have bound more than 3 PI(4,5)P_2_ and almost no SVs form dual bindings at steady state. **B)** Cumulative fusion of SVs after 50 µM step Ca^2+^ at t=0, grouped according to their initial PI(4,5)P_2_ binding state. During the first ∼0.5 ms, release is dominated by SVs having two or three syts bound to PI(4,5)P_2_ prior to the stimulus. The zoom-in shows that the SVs having three or four syts prebound to PI(4,5)P_2_ constitute the majority of release of the first five SVs and therefore determines the release latency. **C)** Cumulative release probability over time of SVs after 50 µM step Ca^2+^ at t=0, grouped according to initial PI(4,5)P_2_ binding state. The dominance of SVs having pre-bound to PI(4,5)P_2_ with two to four syts in panel B is explained by their high release probability compared to SVs with no or only one PI(4,5)P_2_ bound.

**Figure 4 – figure supplement 1:**
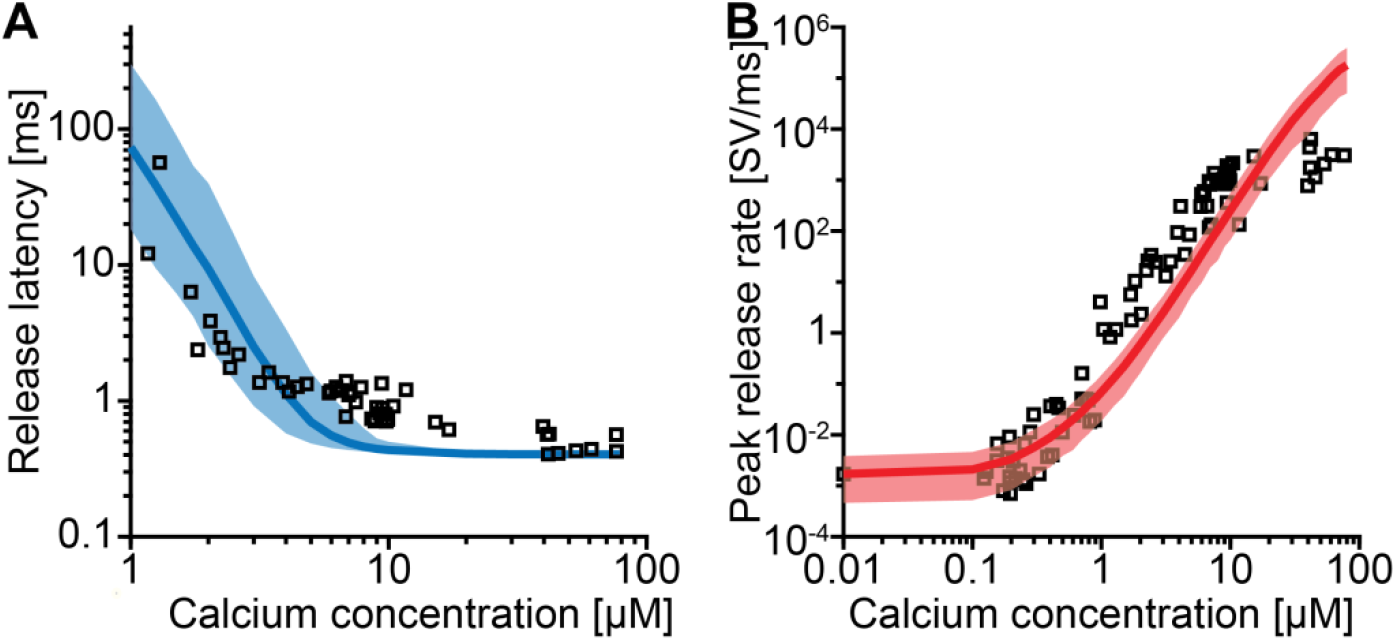
Fitting of the model without allosteric interaction between Ca^2+^ and PI(4,5)P_2_ fails to reproduce the Ca^2+^-dependency of NT release. **A-B)** The best fit results for a model with 3 slots and no allosteric interaction between Ca^2+^ and PI(4,5)P_2_ (A=1). In A the median release latency and the 95% prediction interval of the best fit model are shown. Note that the y-axis range is different from Figure 2A, but the proportions of the ticks is maintained to help comparison. B) The mean maximal fusion rate as a function of [Ca^2+^]_i_ and the corresponding 95% prediction interval. Best fit parameters used to generate these curves are: α=2.174 µM^-2^s^-1^, γ=6.773·10^5^ µM^-1^s^-1^, [PI(4,5)P_2_]=2991 µM, f=3.943·10^5^, d=0.4036 ms. Experimental data points in the figures are replotted from Kochubey and Schneggenburger (2011). Similar to the best fit solutions with too few slots (Figure 2A, left), the fitted *f* value was extremely high, which forced quick SV fusion before the newly formed dual binding dissolved. Thus, when assuming realistic affinities for Ca^2+^ and PI(4,5)P_2_, the allosteric property of the syt C2B domain, which leads to a stabilization of the dual-bound state, is necessary to ensure high Ca^2+^ sensitivity of release.

**Figure 5 – Figure supplement 1:**
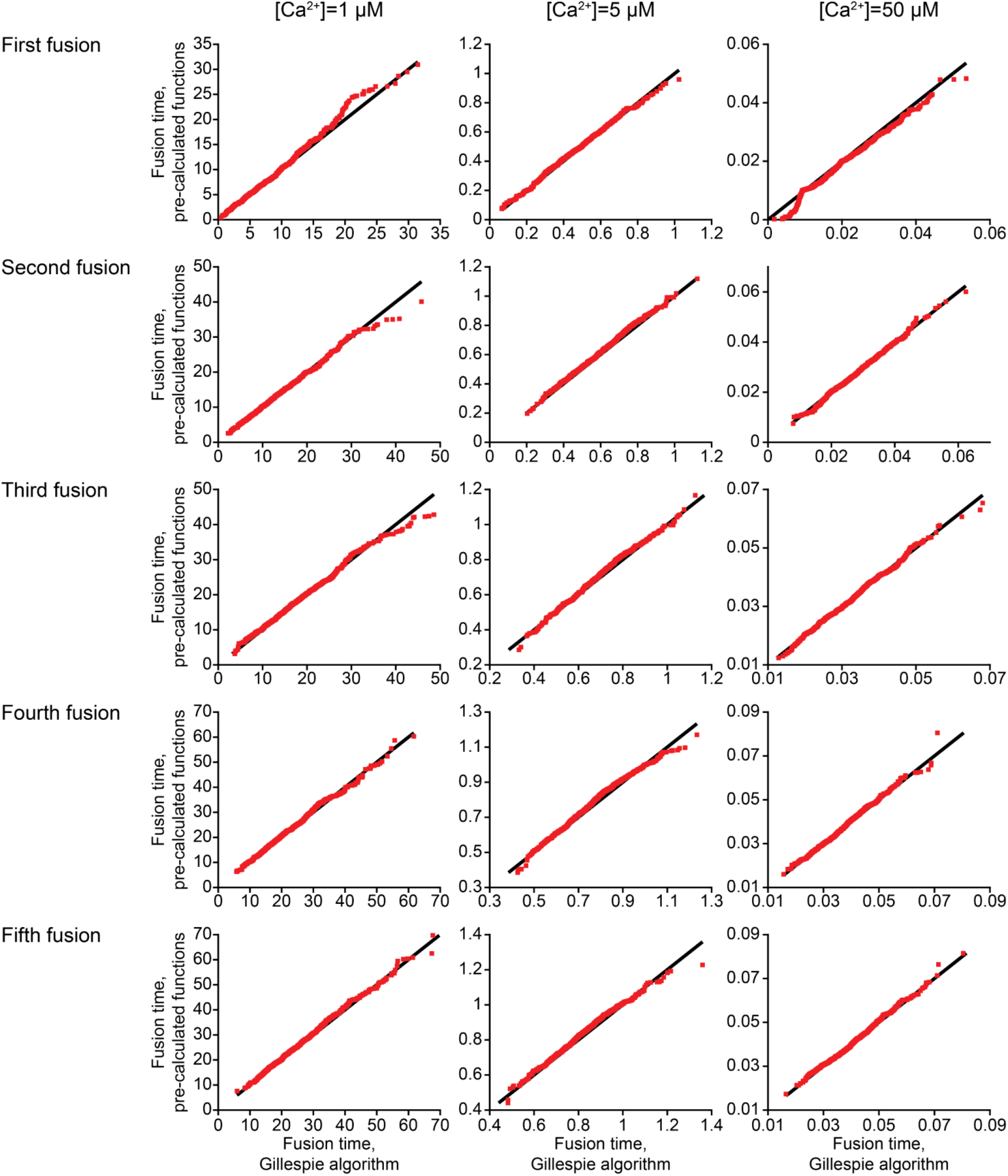
Comparing the two different model implementations. QQ-plots comparing the first five fusion times obtained using stochastic simulations with an implementation based on the closed-form solution of the model and using the Gillespie algorithm for three different [Ca^2+^]_i_ (see <u>Methods</u>). Black line respresent 1:1 correspondence, which would only happen in deterministic simulations. Red squares indicate fusion time simulated with both methods for 1000 repetitions.

**Figure 5 – figure supplement 2.**
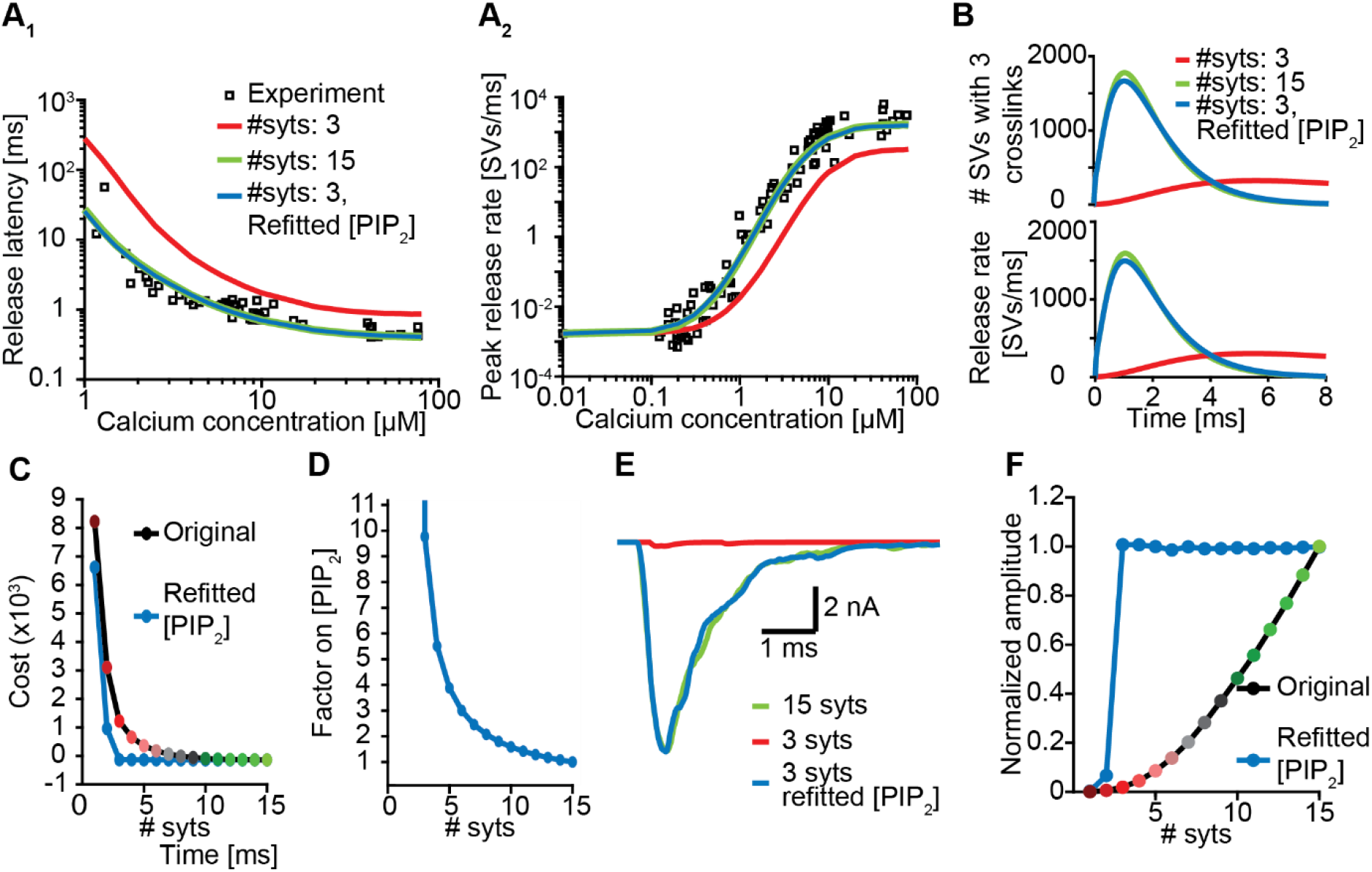
Upregulation of PI(4,5)P_2_ can compensate for loss of syts. **A)** Release latencies (A_1_) and peak release rates (A_2_) for a model with 15 syts (green) and with 3 syts before (red) and after (blue) refitting of [PI(4,5)P_2_]. Experimental data points in panels A are replotted from Kochubey and Schneggenburger (2011).**B)** The average number of SVs with three dual bindings formed (top) and the corresponding release rates (bottom) as a function of time upon stimulation with a Ca^2+^-flash of 50 µM for a model with 15 syts (green), and with 3 syts before (red) and after (blue) refitting of [PI(4,5)P_2_]. **C)** The costs values associated with the Ca^2+^-uncaging data for different levels of syt with the original best fit parameters (black line) and after re-fitting [PI(4,5)P_2_] for each choice of n_syts_ (blue line). **D)** The fold-increase in [PI(4,5)P_2_] obtained by re-fitting the model as a function of the number of syts. **E)** Representative AP-evoked response of a model with 3 syts per SV obtained after increasing [PI(4,5)P_2_] plotted together with representative responses for a model with 3 and 15 syts with the original [PI(4,5)P_2_]. **F)** Average amplitudes of simulated AP-evoked responses at original [PI(4,5)P_2_] and using the increased values.

**Figure 6 – figure supplement 1.**
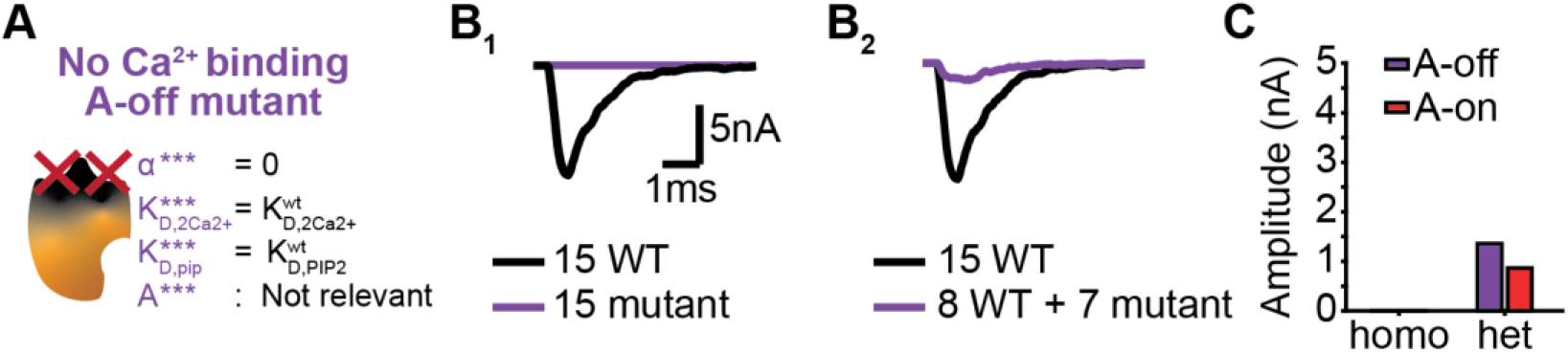
The dominant negative effect of a mutant that is unable to bind Ca^2+^ depend on the mutants PI(4,5)P_2_ affinity. **A)** Schematic illustration of the “*No Ca*^*2+*^*-binding, A-off”* mutant, which is not able to bind Ca^2+^ implying that the allosteric interaction between Ca^2+^ and PI(4,5)P_2_ is always ‘inactive’. **B)** Representative, stochastically simulated AP-responses with homozygous (left, 15 mutants) and heterozygous (right, 7 mutants + 8 WT) of the A-off mutant. For each of the examples representative trace of a condition with 15 WT syts is shown in black. **C)** Comparison of the mean amplitudes of AP-evoked responses (n=200) simulated with “*No Ca*^*2+*^*-binding, A-off”* mutant to the mean amplitudes of the “*No Ca*^*2+*^*-binding, A-on”* mutant (Figure 6).

